# Cationic Polymers Enable Internalization of Negatively Charged Chemical Probe into Bacteria

**DOI:** 10.1101/2023.03.19.533359

**Authors:** Hannah K. Lembke, Adeline Espinasse, Mckenna G. Hanson, Christian J. Grimme, Zhe Tan, Theresa M. Reineke, Erin E. Carlson

## Abstract

The bacterial cell envelope provides a protective barrier that is challenging for small molecules and biomolecules to cross. Given the anionic nature of both Gram-positive and Gramnegative bacterial cell envelopes, negatively charged molecules are particularly difficult to deliver into these organisms. Many strategies have been employed to penetrate bacteria ranging from reagents such as cell-penetrating peptides, enzymes, and metal-chelating compounds, to physical perturbations. While cationic polymers are known antimicrobial agents, their ability to permeabilize bacterial cells without causing high levels of toxicity and cell lysis has not been demonstrated. Here, we evaluate the ability of four cationic polymers, two linear and two micellar (from self-assembled amphiphilic block copolymers), to facilitate the internalization of an anionic ATP-based chemical probe into *Escherichia coli* and *Bacillus subtilis*. Not only did we observe the permeabilization of these organisms, but also found that labeled cells were able to continue to grow and divide. In particular, the micelle-based polymers yielded effective internalization of the negatively charged chemical probe better than their linear analogues.

## Introduction

The bacterial cell envelope is a protective barrier from the environment, and therefore, critical for survival.^1^ Bacteria have evolved highly effective methods to selectively enable desired compounds to cross their cell envelope and to prevent the internalization of others. These structures not only pose major challenges in the development of antibacterial agents, but hinder our ability to efficiently deliver other cargo, such as chemical probes, to the cytosol. Probe delivery requires methods that not only facilitate cell penetration but do so without causing substantial microbial death. Such strategies are crucial for the continued study of bacterial physiology – chemical probes must gain entry across cellular barriers without significant damage.^2–7^ It is also thought that some internalization reagents can play important synergistic roles with antibiotics.^2, 8^

The two major classes of bacteria, Gram-positive and Gram-negative, differ based on the structure of their cell envelope. In Gram-positive bacteria, it is comprised of a cytoplasmic membrane made of phospholipids followed by a thick layer of peptidoglycan containing teichoic acids. The Gram-negative envelope also contains a phospholipid inner membrane (IM), which is in combination with a thin peptidoglycan layer that is surrounded by an additional outer membrane (OM) coated in lipopolysaccharides (LPSs) and surface proteins.^9^ Previous work has demonstrated the transport of compounds to the cytosol in both types of bacteria through a variety of methods including synthetic transporters^10^, hyperporination^4^, use of efflux pump inhibitors^11^, hyper-susceptible mutant strains (such as *E. coli* DC2)^12–13^, electroporation^14^, and membrane permeabilization agents.^2, 15–21^ Of particular importance to these studies, there is significant literature focused on reagents that permeabilize the cell envelope including polyamines^17–18^, the cationic chelator ethylenediaminetetraacetic acid (EDTA),^22^ and cationic peptides,^2, 6, 19–20^ of which the polymyxins are of crucial importance.^21, 23^ Polymyxins are highly toxic, but analogs that act exclusively to permeabilize the cell envelope have also been discovered.^23–25^ For instance, a commercially available polymyxin derivative, polymyxin B nonapeptide (PMBN), is considerably less toxic than its parent compound but retains its permeabilization ability.^15–16^ This property makes it an effective reagent for sensitizing Gram-negative bacteria to antibiotics that are otherwise ineffective. PMBN disrupts the OM of Gram-negative bacteria by interacting with the negatively charged LPS but has a limited effect on bacterial viability. Given the lack of LPS, polymyxins have little-to-no activity against Gram-positive bacteria.^15, 26–27^ The use of cationic polymers to act as antimicrobial agents is well-established due to their cheap cost and easy-to-modulate chemistry.^28–29^ However, the identification of molecules, and in particular cationic polymers, that interact with the cell envelope and promote penetration without causing cell death is a much less developed field.

The delivery of negatively-charged molecules into both prokaryotic and eukaryotic cells is a considerable challenge given the charge-charge repulsion between these species and the cell membrane/envelope structures.^30^ However, this hurdle must be addressed to internalize small molecule probes, such as those based on nucleoside triphosphates (NTPs). Romesberg and coworkers reported an engineered strain of *E. coli* that expresses an NTP transporter, facilitating the internalization of non-natural NTPs.^31^ While successful, importers have limited recognition scope and this strategy cannot be used in wild-type organisms. Kraus and co-workers reported the use of a positively charged cyclodextrin derivative, termed synthetic nucleotide triphosphate transporter (SNTT), to internalize NTPs into eukaryotes and prokaryotes.^10^ SNTT binds the triphosphate moiety masking its charge. It is suspected that the polyarginine tail inserts itself into the cellular envelope to deliver the cargo, followed by NTP displacement with endogenous phosphates. SNTT has been used to deliver cyanine-3-deoxyuridine triphosphate into *E. coli* and *Mycobaterium smegmatis.^10^* Nevertheless, the use of SNTT has only been shown in two organisms and with nucleoside-functionalized NTPs.

An alternative strategy utilizes a prodrug approach in which negatively charged moieties are masked by protecting groups that are cleaved off by endogenous enzymes upon molecule internalization. This method has also been employed in viruses.^32–33^ However, the synthesis of molecules containing masked phosphates remains challenging and incomplete enzymatic cleavage of the prodrug can result in a low intracellular concentration of the active species. Relatedly, in an approach termed ProTide, a monopho(n)sate is converted to its active di- and tri-phosphate form once in the cytosol.^32^ This strategy can also be synthetically challenging and suffer from incomplete *in vivo* conversion to the active molecule, while also precluding the use of nucleotides that are functionalized off of the phosphate(s).

With the goal of devising a highly generalizable internalization method that is applicable to any type of bacteria, we sought inspiration from the rapidly expanding pool of reagents that have proved to be efficient in delivery of negatively-charged genetic materials across cell membrane lipid bilayers and into eukaryotic cells.^34^ Polymer-based delivery systems have been utilized to transfect eukaryotic cells with extracellular genetic materials, such as plasmid DNA (pDNA), RNA, and antisense oligonucleotides, through the formation of complexes held together via electrostatic interactions.^35, 36^ In these systems, the genetic material is condensed and protected within the polymer structure during delivery into cells via an endosome-mediated mechanism. After escaping the endosome, it is translocated to the nucleus and cytosol.^37^ Many of these polymers present nitrogen-containing motifs that are positively charged at physiological pH. As such, the negatively charged phosphate groups of the genetic material interact with the positively charged polymers to form a complex called a polyplex. Reineke and co-workers have extensively studied cationic polymer-based systems for the delivery of payloads into mammalian cells, with a particular focus on the effects of polymer architecture. One key discovery has been the importance of complex architectures such as micelles (termed micelleplex when complexed with genetic material) for better delivery.^38–42^

The cationic nature of these materials suggests that they may also interact favorably with bacterial cells^2, 20, 28, 43–44^ In addition, the ease of synthesis of these types of molecules could make them a promising new class of bacterial delivery reagents. While eukaryotic membranes and prokaryotic cell envelopes differ, the ability of these polymers to complex with negatively charged genetic materials makes them promising candidates for the delivery of other negatively charged molecules such as the adenosine triphosphate (ATP)-based probe, BODIPY-FL-ATPγS.^45^ We have previously reported the use of this molecule to study the activity of bacterial histidine kinases, sensory proteins involved in signaling. Herein, we investigate the ability of four polymer architectures to complex and deliver ATP-based probes into both Gram-negative and Grampositive bacteria, resulting in the identification of polymeric materials that enable efficient probe delivery. These novel internalization reagents could be applied to the study of various NTP-related processes in live bacteria, which is currently an extreme challenge. They may also find general utility for the delivery of myriad anionic molecules into the bacterial cytoplasm, ranging from genetic material to antibacterial agents.

## Results

### Tools for chemical probe internalization

#### Polymer selection

We sought to investigate polymers that have recently shown great promise for the delivery of challenging payloads [i.e., pDNA and antisense oligonucleotides (ASO)^38, 40^] into mammalian cells, including two linear polymers and two polymers that can self-assemble into micelles. The hydrophilic poly(ethylene glycol) (PEO or **O**), cationic poly(2-(dimethylamino)ethylmethacrylate (PDMAEMA or **D**), and hydrophobic poly(*n*-butyl methacrylate) (PnBMA or **B**) were utilized as building blocks to prepare polymeric micelles or linear polymers. The **O** block enhances the hydrophilicity of the polymer. The amine-containing **D** block is cationic under physiological conditions due to its optimal pKa (~8.5)^38^, which, in eukaryotic cell applications, interacts through electrostatic forces with polyanions, such as nucleic acids or polysaccharides on the cell membrane.^38^–^40^ The hydrophobic **B** block enables the formation of the micelles by self-assembly, acting as their core and exposing the other blocks (**O** or **OD**) to the environment. The micelles were formed using the PDMAEMA-*b*-PnBMA (**DB**) amphiphilic diblock copolymer or PEO-*b*-PDMAEMA-*b*-PnBMA (**ODB**) triblock copolymer, yielding micelles with hydrodynamic radii of 28 nm and 34 nm, respectively. The linear polymers were built with hydrophilic building blocks only and formed PEO-*b*-PDMAEMA (**OD**) diblock polymer or PDMAEMA (**D)** homopolymer (**Fig. 1A**).

**Figure 1:**
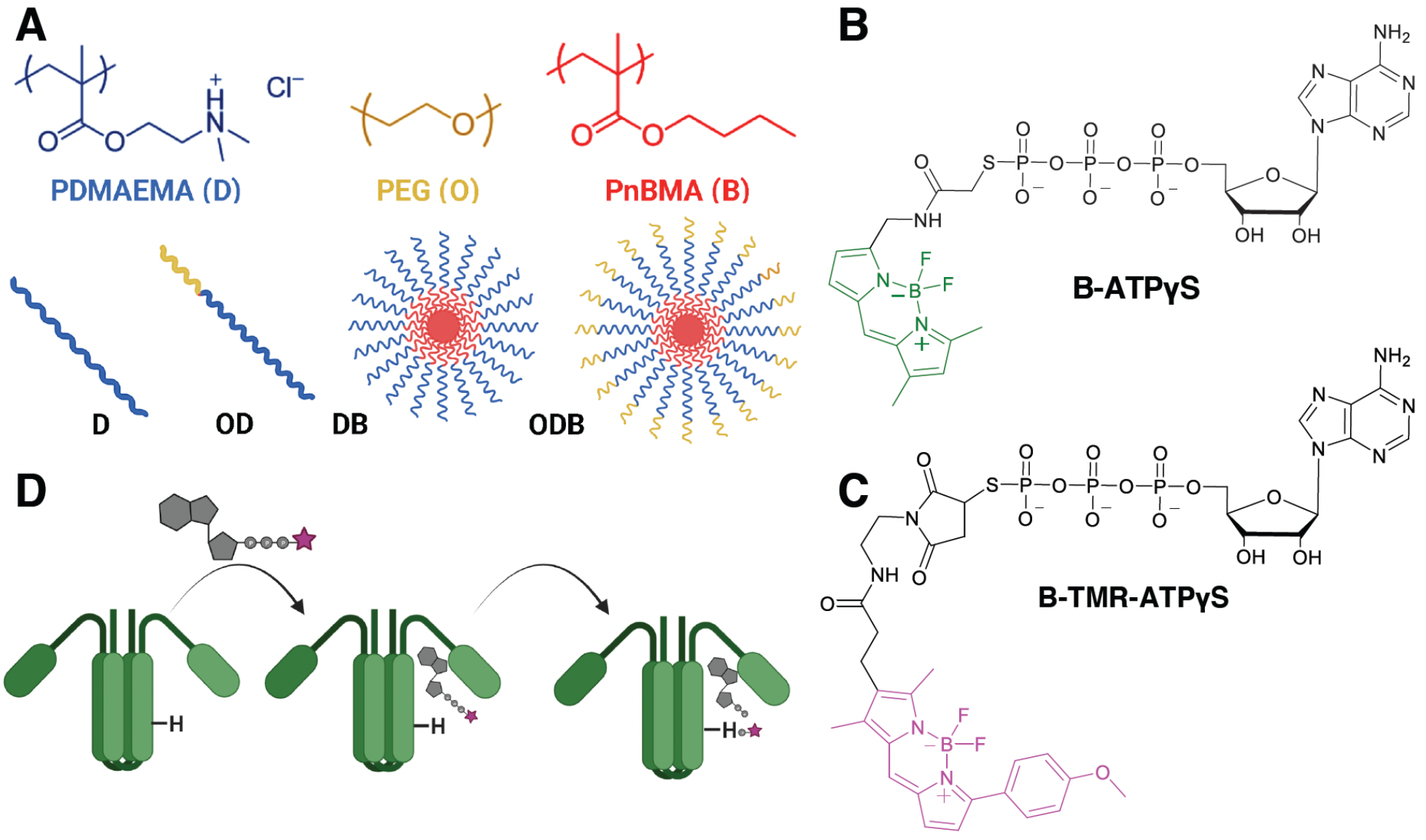
Probe and Polymer Structures. **A.** Component blocks for the polymers and cartoon depictions of their overall architecture. **B.** Structure of **B-ATPγS** activity-based probe. **C.** Structure of **B-TMR-ATPγS** activity-based probe. **D.** Cartoon depiction of fluorophore transfer from ATP-based probe to model HK, HK853.

Our prior work demonstrated that all four vehicles, linear polymers (**D** and **OD**) and micelles (**DB** and **ODB**), complex with pDNA^40^ and ASO^38^ and deliver these biomolecules into eukaryotic cells. Interestingly, there were substantial differences in delivery efficacy based on the polymer architecture, with micelles performing significantly better than linear polymers. It is thought that the higher concentrations of amines condensed and packed together within the micelle may stimulate both internalization and endosomal escape in mammalian cells. Indeed, this system was able to deliver challenging payloads to mammalian cells, such as CRISPR-Cas9 cargo, even outperforming industry standards.^39^ Here, we explore the utility of this suite of polymers for delivery into bacteria. We hypothesized that the negatively-charged phosphate tail of an ATP derivative could interact with the cationic polymers, promoting its internalization into bacteria.

#### Chemical Probe Design

To enable rapid assessment of the internalization and subsequent cytoplasmic accessibility of an ATP-derivative, we sought to use a functionalized analog, BODIPY-FL-ATPγS (B-ATPγS; **Fig. 1B**), that we previously demonstrated will label catalytically-active bacterial histidine kinase (HK) proteins.^45, 46^ This activity-based probe (ABP) results in the transfer of a fluorescent phosphate moiety directly to the targeted proteins, promoting rapid visualization in gel-based assays (**Fig. 1D**). Initial internalization studies in *E. coli* with B-ATPγS suffered from high background fluorescence that overlapped with the BODIPY-FL spectra, likely due to intrinsically-fluorescent flavoproteins ^47–48^ (**Fig. S1**). To overcome this challenge, we synthesized a related analog with a red-shifted fluorophore, BODIPY-TAMRA (λ_ex_/λ_em_ 542 nm/568 nm). Insertion of the anisole moiety causes a shift in excitation and emission of the fluorophore, resolving overlap with the inherently fluorescent components of the bacterial proteomes. BODIPY-TAMRA-ATPγS (**B-TMR-ATPγS**) was synthesized in one step via a Michael-type reaction from ATPγS with BODIPY-TAMRA-maleimide (**Fig. 1C**, **Scheme S1**). **B-TMR-ATPγS** was verified as an ABP by labeling of a purified HK, a constitutively-active construct of HK853 from *Thermotoga maritima*^45, 48^ (**Fig. S2**).

#### Permeabilization of E. coli with cationic polymers

To evaluate the ability of cationic polymers to facilitate entry of **B-TMR-ATPγS** through the bacterial cell envelope, we investigated internalization into a strain of *E. coli* (BL21) harboring an inducible vector for overexpression of HK853 (constitutively-active). We evaluated probe internalization using our previously established gel-based assay^45–46^ to visualize HK853 that is fluorescently labeled by **B-TMR-ATPγS** when this probe successfully reaches the cytoplasm. Our first objective was to optimize the concentrations of both polymer and probe, initially with respect to one another, and to ultimately decrease the overall concentrations to minimize background labeling and cell perturbations. We tested a range of nitrogen/phosphorous (N/P) molar formulation ratios, calculated from the number of protonatable nitrogen atoms in the polymer **D** block to the phosphate groups in **B-TMR-ATPγS**. At each N/P ratio, **B-TMR-ATPγS** (33 μM) was first incubated with the polymer to promote complexation (30 min), and this mixture was then added to HK853-expressing *E. coli* (30 min). Following cell lysis, the proteome was separated via SDS-PAGE and the labeled proteins visualized by fluorescence scanning. A 5/1 N/P ratio yielded optimal protein labeling with the lowest possible amount of polymer, which was also previously found to be optimal for pDNA delivery in eukaryotic cells (**Fig. S3**)^40^. Next, we determined that comparable protein labeling could still be achieved with a lower probe concentration (20 μM) when the 5/1 N/P ratio was maintained (**Fig. S4**).

Using these optimized conditions, we compared the efficacy of our small polymer library, as well as two known internalization agents, PMBN and SNTT (**Table S1** and **Fig. S5**). We found that the micelle polymers yielded the highest HK853 labeling (assigned as 100%), followed by the linear polymers [94% (**OD**) and 72% (**D**) relative to micelle polymers]. The commercially available reagents, SNTT and PMBN, trailed at 22% and 37% relative HK853 labeling, respectively. We also found that a control polymer containing only the **O** block (i.e., PEG functionality without amines) was insufficient for protein labeling (**Fig. S6**). The hydrophobic **B** block alone was insoluble in media. Finally, we confirmed that although the hydrophobic nature of a fluorophore may promote non-specific protein labeling,^49–50^ this was not observed in our experiments (treatment with **ODB** or **DB** and BODIPY fluorophore; **Fig. S7**).

#### Bacterial damage following polymer exposure

Permeabilization reagents are bactericidal if they cause too much damage.^24^ Promisingly, initial evaluation of the antibacterial properties of these polymers indicated a lack of cell killing with both Gram-positive and Gram-negative bacteria (**Fig. S8**). However, we sought more specific information to determine if our ability to label HK853 was due to internalization of the probe or was the result of wide-spread cellular damage that exposed HK853 and other cytoplasmic proteins to **B-TMR-ATPγS**. Pr differentiate live and d DNA.^51^ PI uptake resu (negative control) or co to **B-TMR-ATPγS**. Propidium iodide (PI), a membrane-impermeable DNA dye, is used to differentiate live and dead cells by measurement of its entry through the cell envelope to reach DNA.^51^ PI uptake resulting from polymer treatment was evaluated in comparison to untreated (negative control) or colistin-treated (positive control) cells. Colistin is an antibiotic that causes permeabilization of the OM in Gram-negative bacteria.^52^ We found that the micellular structures and SNTT caused significantly less damage to the *E. coli* envelope than either the linear polymers or PMBN (**Fig. 2A**).

**Figure 2:**
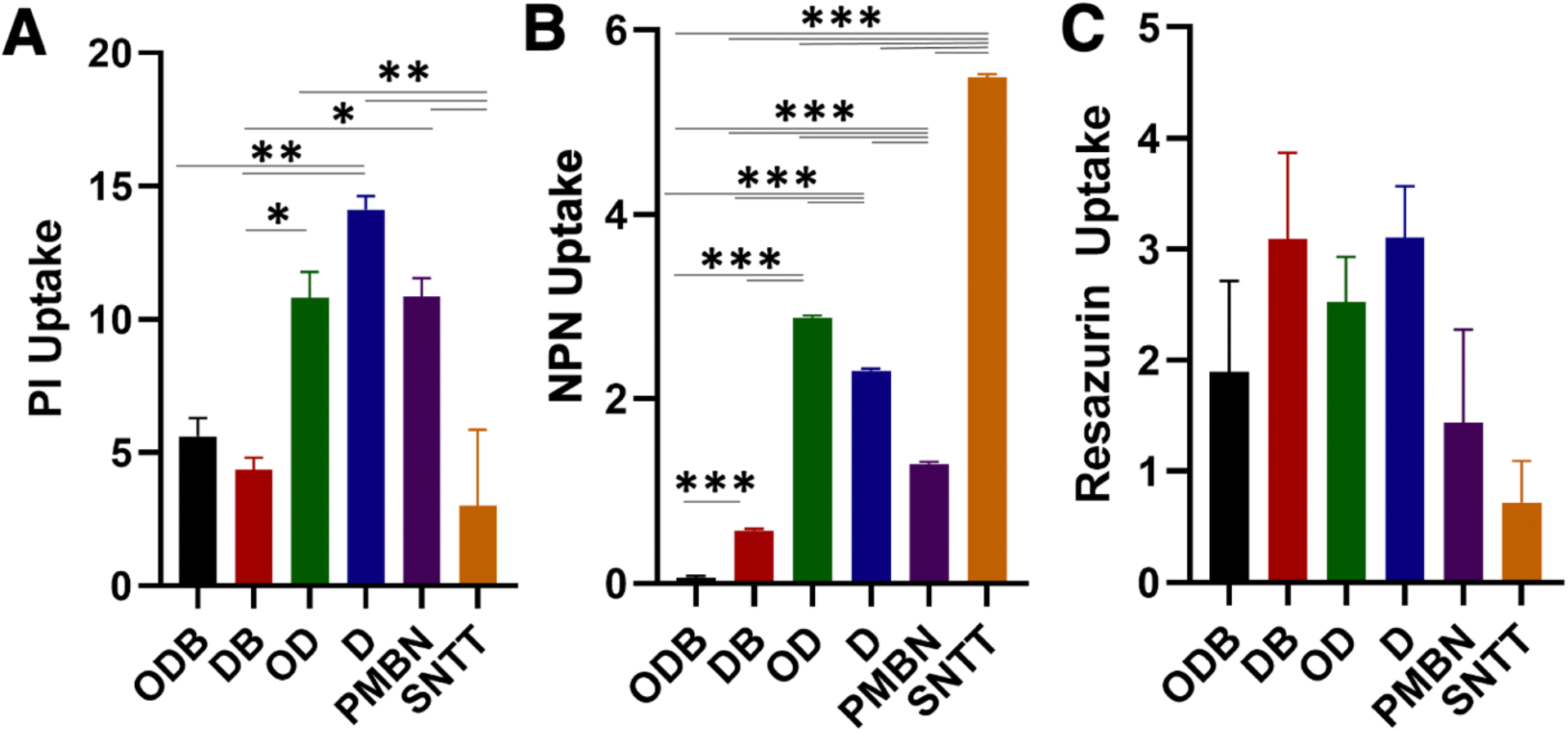
Cellular damage following treatment with cationic polymers. **A.** Cell envelope permeabilization. Propidium iodide uptake following treatment with each polymer (5/1 N/P calculated from 20 μM **B-TMR ATPγS**), PMBN (200 μM), and SNTT (40 μM) compared to untreated control after 30 min. **B.** Outer membrane permeabilization. Comparison of NPN uptake following treatment with each polymer (5/1 N/P calculated from 20 μM **B-TMR ATPγS**), PMBN (200 μM), and SNTT (40 μM) compared to untreated control after 15 min. **C.** Metabolic activity. Resazurin treatment after incubation with each polymer (5/1 N/P calculated from 20 μM **B-TMR ATPγS**), PMBN (200 μM), and SNTT (40 μM) compared to untreated control after 10 min. Error bars represent standard deviations from two biological replicates. Uptake calculations in the methods section. All statistical analysis performed in GraphPad Prism using One-Way ANOVA using Tukey’s multiple corrections. * Indicates p-value <0.05, ** p-value <0.005, *** p-value <0.001.

Next, we specifically assessed OM damage using 1-*N*-phenylnaphthylamine (NPN), which becomes fluorescent in hydrophobic environments.^7^ As such, when NPN interacts with the IM, it becomes highly fluorescent and enables us to evaluate damage strictly to the OM. We hypothesized that there may be interactions between the positively-charged **D** block in the polymers and the negatively-charged LPS of the Gram-negative OM, potentially resulting in the formation of pores through the membrane.^53^ Our data indicate that the linear polymers cause OM damage similar to PMBN, whereas the micelle polymers yield substantially less (**Fig. 2B**). SNTT treatment, which is believed to involve the polyarginine tail leafing through the OM,^10^ resulted in the largest perturbation. These data may indicate differences in the membrane disruption mechanisms of the investigated reagents.

Finally, we evaluated how the polymers affected the metabolic activity of treated cells as a measure of cell viability. Resazurin (blue and weakly fluorescent) is reduced intracellularly to resorufin (pink and highly fluorescent) by electron receptors such as NADH dehydrogenase in live cells.^54–55^ Reductase activity correlates with organism respiration and thus correlates with cell viability. We observed that all treatments, with the exception of SNTT, yielded higher fluorescence signal than the untreated control (**Fig. 2C**; polymers are not intrinsically fluorescent, **Fig. S9**), which indicate low toxicity of the polymers. We anticipate that this increased fluorescence is likely due to the enhanced permeability of the cellular envelope, enabling a higher effective concentration of resazurin to enter the treated cells. This is supported by our observation that colistin, a well-known bactericidal agent, and PMBN also cause increased fluorescent signal (**Fig. S10**).^56^ Combined, these assays indicate that the micelle **DB** polymer shows the best combination of membrane permeabilization and cell viability.

While the PI and resazurin assays give insight about the overall viability of the cell population and the gel-based assay indicated internalization of **B-TMR-ATPγS**, we sought to further investigate the health of individual labeled cells. These data are needed to confirm that cells penetrated with the probe are still able to grow, as population-level experiments do not exclude the possibility that labeled cells die and unlabeled cells are those that appear to be viable following treatment. We performed time-lapse microscopy experiments to track cells treated with **B-TMR-ATPγS** and the micelle **DB** polymer, which showed the most promise in previous assays, or the related linear **D** polymer. We found that cells with internalized probe, as judged by the presence of fluorescent signal, were able to grow and propagate (**Fig. 3** and **S11**). We also observed that a proportion of the labeled cells lysed or became filamented, a known response to cell membrane damage.^57^ This phenomenon was much more pronounced following treatment with **D** than **DB**, which is consistent with the results of our membrane damage assays showing that **D** polymers lead to more damage. As expected, treatment with **B-TMR-ATPγS** alone yielded no fluorescent cells and normal cell growth curves (**Fig. S12**), while treatment with **DB** or **D** alone caused filamentation (**Fig. S13** and **S14**). Filamentation was also seen in cells treated with PMBN suggesting that many permeabilization agents may induce this phenotype (**Fig. S15** and **S16**). Cells treated with SNTT showed a high propensity for lysis (**Fig. S17** and **S18**).

**Figure 3:**
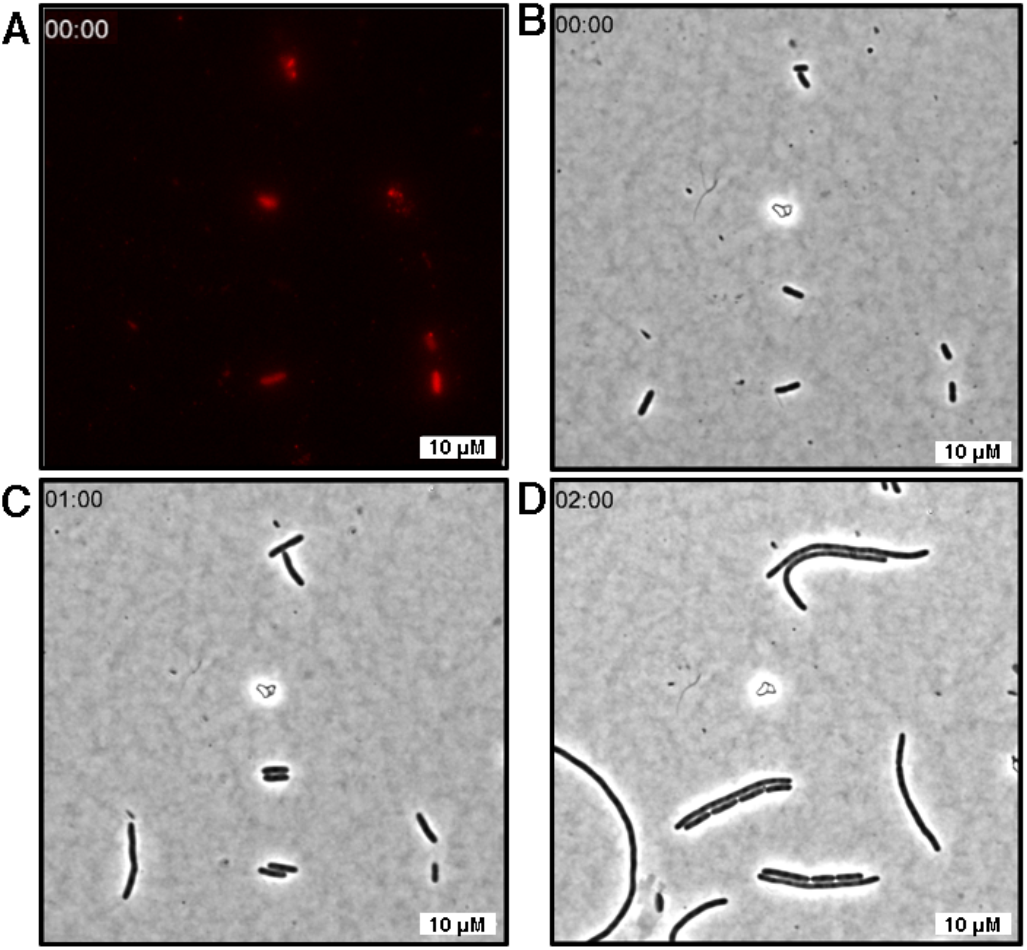
Time-lapse imaging of live *E. coli* treated with DB and B-TMR-ATPγS. **A**. Zero timepoint (t=0) demonstrates labeling with **B-TMR-ATPγS** (20 μM with 0.073 mg/mL **DB** polymer in TES buffer; imaged in TRITC channel; λex/λem 542 nm/568 nm). Fluorescence photobleached during the course of the experiment making fluorescent imaging impossible at later timepoints. **B.** Bright field (BF) channel at t=0. **C.** BF channel at t=1 h. **D.** BF channel at t=2 h. Scale bars are 10 μm. Representative images taken from a single time-lapse experiment out of three biological replicates. Movies included in Supporting Information.

#### B-TMR-ATPγS and Polymer Complexation

While the polymers enabled **B-TMR-ATPγS** to effectively penetrate the cellular envelope of *E. coli*, it was unclear if they complexed with **B-TMR-ATPγS** to facilitate entry into the cell or serve exclusively as a permeabilization agent. While prior work has demonstrated clear interactions between this suite of polymers and pDNA and ASO^38, 40^, complexation with a substantially smaller molecule has not been investigated. First, we compared the level of HK853 labeling when the polymers were either used to pre-treat the cells or when they were added at the same time as **B-TMR-ATPγS**. We reasoned that if probe internalization only required permeabilization by the polymers, and not complexation between the polymer and **B-TMR-ATPγS**, we should still see substantial protein labeling when the cells were first treated with polymer and then with probe. Gel-based assessment revealed decreased band intensities in polymer pre-treatment samples (polymer followed by brief wash and probe incubation; **Table S2**). Coomassie staining to evaluate protein loading confirmed that the decreased fluorescence signal was not due to overall lower protein abundance (**Fig. S19**). While the labeling intensity was diminished, the same protein banding pattern was observed, suggesting that complexation (polymer and probe preincubation) is not required for probe entry into the cytosol (**Fig. 4**). Entry into the cytosol without polymer complexation has also been demonstrated in eukaryotic cells for the delivery of pDNA^58^. We also evaluated pre-treatment versus co-treatment with PMBN and SNTT. We observed no appreciable difference in the intensity of the labeled proteins (**Fig. S20**), further indicating that our polymer library is likely promoting probe entry through a mechanism that is distinct from these reagents.

**Figure 4:**
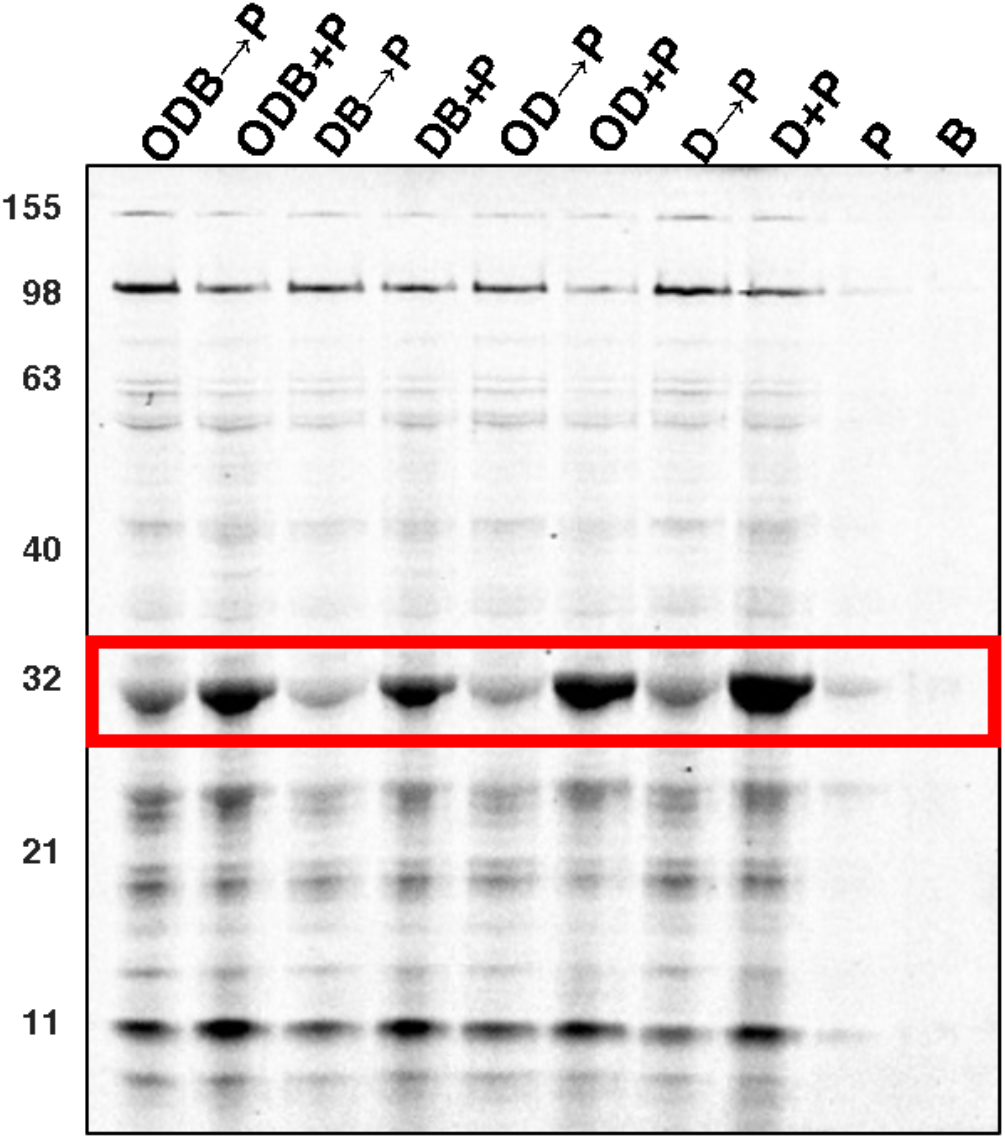
Polymer pre-treatment versus co-treatment of *E. coli*. Arrow (→) indicates pretreatment (30 min) with polymer (5/1 N/P) prior to incubation with **B-TMR-ATPγS** (P, 20 μM, 30 min). Pre-treated samples were washed once (100 μL TES buffer) between incubations. Plus (+) indicates co-treatment with the polymer and probe at the same concentrations (30 min). B is buffer alone. Representative gel from two biological replicates. Red box indicates HK853. Molecular weight ladder in kDa.

To further evaluate if detectable interactions occur between our anionic probe and the cationic polymers, we employed dynamic light scattering (DLS) and electrophoretic mobility shift assays, which are routinely used to study polymer/nucleic acid complexes. While most investigations focus on large oligonucleotides, interactions have been characterized between cationic polymers and ASOs (~20 negative charges), indicating that smaller polyion species can favor complexation.^59^ We performed DLS studies similar to those previously reported with **ODB**, **DB**, **OD**, and **D** and pDNA^40, 58^ and observed no significant change in the size of the micelles upon incubation with **B-TMR-ATPγS** (**Fig. S21**). The linear polymer/probe mixtures yielded species that were too small to reliably measure. However, there was no significant difference in the normalized scattering intensity when the probe was added, indicating that there was no change in size upon probe addition (**Table S3**). Given the lack of size changes observed upon probe/polymer incubation, we concluded that no significant interactions were taking place. However, this result may not be conclusive since a small probe might not induce an observable size change. As such, we obtained additional evidence with an electrophoretic mobility shift assay. Probe complexation would result in a molecular weight change and thus, a band shift on the gel. No shift was seen with any polymer, but it is still possible that the changes were too small to be detected with this method (**Fig. S22**). Finally, we employed a method with superior sensitivity, ^31^P NMR, which provides a direct readout of the phosphorus environment. As a proxy for our supply-limited **B-TMR ATPγS** probe, ATP was used as it should have similar electrostatic interactions with the cationic polymers. Spectra were collected after incubation of ATP and the **D** polymer in TES buffer (30 min; **Figure S23**). The micelle polymers were formulated at concentrations that are too low for use in these experiments. We observed no change in comparison to our ATP control (**Figure S24**) providing further evidence that no interaction occurs between the polymers and ATP, or its derivatives. Thus, we conclude that probe/polymer complexation is not required for cellular entry.

#### Permeabilization of Gram-positive B. subtilis

While it is generally accepted that Gram-positive organisms are easier to penetrate than their Gram-negative relatives, they present unique challenges as their cell envelope has a distinct architecture, including a thick cell wall and teichoic acids.^1, 30, 60^ We sought to determine if the Gram-positive cell envelope could also be penetrated with our suite of polymers using the model bacterium *Bacillus subtilis*. As with *E. coli*, co-treatment of *B. subtilis* with each polymer and **B-TMR-ATPγS** showed substantial protein labeling that was dependent upon the presence of the polymers (**Fig. 5A**; Coomassie stain in **Fig. S25**). We also evaluated cell damage and viability using a subset of the previous assays. PI uptake remained near 0.5 in *B. subtilis* for treatment with **ODB** and **DB**, compared to near double this following treatment with **OD** and **D**, indicating that the micelle polymers may cause less cell death, although the differences were not statistically significant (**Figure 5B**). Assessment with the resazurin assay showed that cell viability following polymer treatments remains above 50% in all cases (**Fig. 5C**). Overall, the differences in cellular damage caused by the four polymers that we had observed in *E. coli* were much less pronounced in this Gram-positive organism.

**Figure 5:**
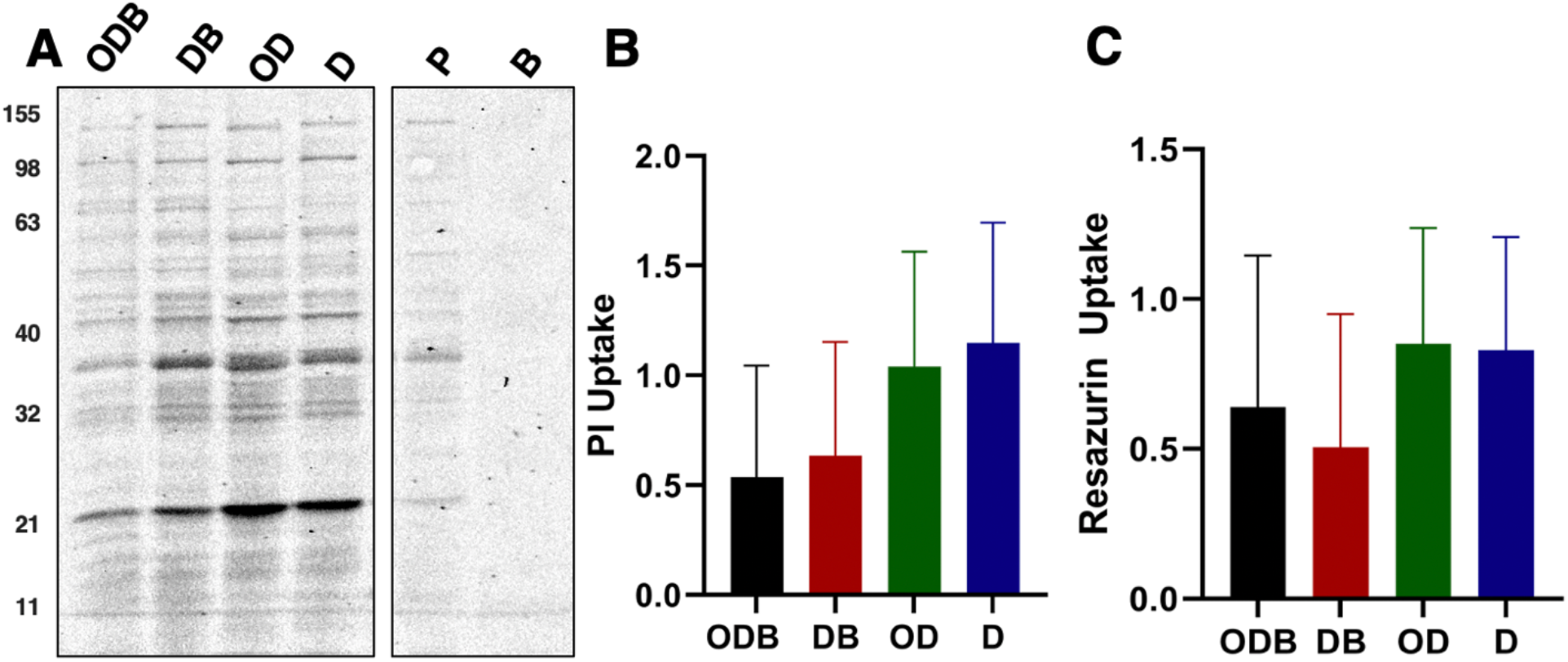
*B. subtilis* labeling and viability following polymer treatment. **A.** Labeling profile of *B. subtilis* upon treatment with polymer (5/1 N/P ratio) and **B-TMR-ATPγS** (20 μM) for 30 min. Probe only (P, 20 μM) and TES buffer (B) lanes included as controls. Representative gel from three biological replicates. Molecular weight ladder in kDa. Coomassie stained gel in **Figure S25**. **B.** Cell envelope permeabilization. PI uptake after treatment with each polymer (5/1 N/P, 20 μM **B-TMR-ATPγS**, 30 min) compared to a lysozyme control (2 mg/mL). Error bars represent standard deviations from two biological replicates. **C.** Metabolic activity. Resazurin uptake following treatment with each polymer (5/1 N/P, 20 μM **B-TMR-ATPγS**, 30 min) compared to resazurin uptake in untreated cells. Error bars represent standard deviations from two biological replicates. Statistical analysis performed in GraphPad Prism using one-way ANOVA indicated no significant differences.

## Discussion

Our studies show that all four polymers enabled the labeling of cytosolic proteins with the anionic chemical probe **B-TMR-ATPγS** and yielded superior labeling compared to commercially available reagents PMBN or SNTT (**Table S4** provides labeling values normalized to protein content). Interestingly, our data indicate that probe internalization is likely occurring through a different mechanism than either the polymers facilitate the internalization of pDNA into eukaryotic cells or by which SNTT promotes molecule entry. Indeed, we see no evidence of a stable complex formed between **B-TMR-ATPγS** and the polymers (e.g., DLS, electrophoretic mobility shift, NMR), but instead anticipate that they are acting as permeabilization reagents on the cells.

We anticipate that our polymer library promotes entry most similarly to the polymyxins. While the mechanism of action of the polymyxins is not fully understood, it is thought that initial electrostatic interactions with the negatively charged phosphate groups of the LPS, followed by displacement of native metal cations (Mg^2+^ and Ca^2+^), is a crucial step. Since all evaluated polymers contain a positively-charged **D** block, we anticipate that they may initially interact similarly with *E. coli*.^25^ As such, we investigated the damage that these polymers caused to the cellular envelope of *E. coli* and found that the best-performing polymers (micelle architecture, **ODB** and **DB**) cause less damage to the overall cell envelope (PI uptake) or the outer membrane (NPN uptake) than PMBN. This is an intriguing result given that these polymers also enable superior protein labeling (**Table S4**).

Deeper analysis of the OM perturbation data indicates that **ODB** caused less damage than **DB**. This may be due to a specific interaction between the OM and the outer-most **O** block of the **ODB** polymer, which is composed of a PEG chain, a well-studied hydrophilic polymer with low toxicity.^58^ Perhaps this extra hydrophilic block provides additional space between the LPS and the charged **D** block of the micelle polymer, which could be beneficial as direct interactions may result in a larger perturbation to the OM. This is further supported by the fact that the highest degree of damage is seen upon treatment with the **D** block alone, followed by the **OD** polymer. Since the **OD** polymer is linear (unlike **ODB** where the **O** block creates a corona blocking the **D** block), the **O** block may not provide “protection” to the LPS. Finally, while all polymers were used at the same N/P ratio, we anticipate that the linear polymers may interact with a larger section of the bacterial cell envelope since it is theoretically possible for them to uncoil on this surface, whereas the micelle polymers could only interact through the portion of **D** block that radially extends from the **B** block center. We expect that less overall interaction with the cell envelope may result in the decreased cellular damage that we observed. However, it is not immediately clear how this would also result in superior probe entry/protein labeling (**Table S4**). Future mechanistic studies will aim to better understand interactions between the negatively charged probe, the cationic polymers, and the bacterial cell surface.

The second critical characteristic of the examined polymer library is its minimal bactericidal or bacteriostatic activity. This was demonstrated through a plate-based growth assay as well as the resazurin assay, which showed high cell metabolic activity. Most importantly, livecell imaging provided conclusive evidence that **B-TMR-ATPγS**-labeled cells are still able to propagate. These data also showed that like PMBN, polymer exposure results in some cell filamentation. This phenotype can lead to abnormal growth curves in response to stress but enables cells to continue living, which is crucial for the subsequent study of the labeled population. Together, these results are particularly exciting as many cationic polymers are antibacterial agents, and there is a large body of literature expanding on their potential as antibiotics.^29^ Our data provide one of the first demonstrations of cationic polymers serving as permeabilization agents without causing substantial bacterial death. To our knowledge, the only example of related work was reported by de Souza and coworkers where they showcased the ability of cationic copolymers to promote the transformation of pDNA into a non-competent *E. coli* DH5α strain.^43^ Transformation with these polymers yielded CFU in the range of 10^3^/μg of DNA, whereas a normal transformation ranges in 10^6^ – 10^7^ CFU/μg of DNA.^43, 61^

Finally, our data indicate that the investigated polymers may provide a generalizable strategy for bacterial permeabilization as they also promoted probe entry into the Gram-positive organism, *B. subtilis*. These cells remained more viable and showed less damage than those treated with lysozyme, an enzyme used to degrade peptidoglycan in the cell wall. Interestingly, linear polymers **D** and **OD** seemingly caused less damage in *B. subtilis* than in *E. coli*, potentially indicating that the thicker cell wall provides a stronger barrier to the linear polymers than the OM of Gram-negative organisms.^6^

## Conclusion

The cationic polymers studied herein showed efficacy in permeabilizing the membrane of both Gram-negative *E. coli* and Gram-positive *B. subtilis* while maintaining a high level of cell viability. When compared to other permeabilization reagents like PMBN, the protein labeling profiles obtained upon treatment with our polymer library revealed better permeabilization as judged through higher intensity bands. We also found that cells were less damaged and had a higher level of metabolic activity. In sum, we have identified a small suite of polymers as promising reagents for the internalization of chemical probes into the bacterial cytosol. We anticipate that these polymers, as well as the large library of polymers that have already been studied for eukaryotic gene delivery, will find utility for the entry of NTP derivatives and other chemical probes into bacteria. In particular, micelle architectures appear to provide an ideal mix of penetration and relatively limited cell damage, opening up many possible avenues for advancing our understanding of the microbial world.

## Acknowledgements

We gratefully acknowledge funding as follows: National Institutes of Health Biotechnology Training Grant 5T32GM008347 (HKL and AE), University of Minnesota Department of Chemistry Excellence Fellowship (HKL), and National Science Foundation CHE2003208 (EEC). We also thank Genentech for partial funding of this project. M.G.H. acknowledges support from the National Science Foundation Graduate Research Fellowship Program (DGE1839286) and the University of Minnesota Doctoral Dissertation Fellowship. This work was also supported by the National Science Foundation (NSF) through the University of Minnesota MRSEC under award DMR2011401. Any opinions, findings, and conclusions or recommendations expressed in this material are those of the authors and do not necessarily reflect the views of the National Science Foundation. BioRender was used for figure generation.

## Author Contribution Statement

HKL: Generated data for figures within the main manuscript and SI, manuscript writing, project direction; AE: Project design and preliminary studies, synthesis of probe and generated data for SI, manuscript writing, project direction; MGH: Design and execution of DLS, intellectual and writing contributions; CJG: Polymer synthesis, intellectual and writing contributions; ZT: Initial project design and polymer synthesis.

## Experimental section

The four polymers used in this study were synthesized and characterized in prior work by Jiang et al.^62^ **B-TMR-ATPγS** synthesis, complexation studies of probe and polymers with DLS, and electrophoretic mobility shift assay, cell growth assay, as well as general experimental details are provided in the Supplementary Information.

### Cell Culture

The overexpression strain of *E. coli* BL21 containing the pHisǁ HK853 plasmid was streaked onto LB agar plates containing 100 μg/mL ampicillin from a frozen glycerol stock and grown overnight at 37 °C. From these plates, a single colony was inoculated into fresh Lennox Broth (LB) media containing 100 μg/mL ampicillin and grown overnight at 37 °C, shaking at 220 RPM. A 1:10 dilution of the overnight culture was performed in fresh LB media containing 100 μg/mL ampicillin and grown until OD_600_ ~0.5-0.6 (Genesys 30 Visible Spectrophotometer). Cells were induced with 1 mM IPTG for 30 min at 37 °C, shaking at 220 RPM. Cells were treated in accordance with each assay. *B. subtilis* 3610 was streaked onto LB agar plates from frozen glycerol stocks and grown overnight at 37 °C. From these plates, a single colony was inoculated into fresh LB media and grown overnight at 37 °C, shaking at 220 RPM. A 1:10 dilution of the overnight culture was performed in fresh LB media and grown until OD_600_ ~0.5. Cells were treated as described for each assay.

### General Procedure for Gel-Based Assays

Prior to addition to cells, 20 μM **B-TMR-ATPγS** was incubated on a rotator in the dark at room temperature with 0.1 mg/mL **ODB**, 0.073 mg/mL **DB**, 0.068 mg/mL **OD**, or 0.05 mg/mL **D**, 200 μM PMBN, or 40 μM SNTT in 30 mM TES buffer (pH = 7.4) for a total volume of 45 μL for 30 min. Amount of polymer calculated based on 5/1 N/P ratio of amines in the polymer to phosphates in 20 μM **B-TMR-ATPγS** probe. Bacteria were spun down (2,000x*g*, 1 min) from the cultures (500 μL for *E. coli;* 1 mL for *B. subtilis*), supernatant removed, and each sample was resuspended with the designated probe/polymer solution or probe/permeabilization reagent solution (40 μL) and then placed in an incubator/shaker (30 min, 37 °C, 220 RPM) in the dark. Cells were centrifuged (2,000x*g*,1 min), supernatant removed, and cells washed once with TES buffer (100 μL). *E. coli* cells were resuspended in TES buffer (40 μL) and 10% (w/v) aqueous SDS solution (4 μL) was added. Cells were lysed on ice by sonication using Hielscher vial tweeter UP200St at 90%A, 70%C for 30 sec on/30 sec off for a total time of 3 min. *B. subtilis* cells were resuspended (40 μL) in a 10 mg/mL lysozyme aqueous solution and incubated in an incubator/shaker (37 °C, 220 RPM, 30 min) in the dark. Following this incubation, 10% w/v aqueous SDS solution (4 μL) was added, and cells were lysed following the aforementioned conditions. Cell lysates were centrifuged (20 min, 21,000x*g*) and the supernatant (20 μL) removed and added to 4X gel loading buffer (8.6 μL). Samples (12 μL) were loaded onto a 10% polyacrylamide gel for SDS-PAGE analysis. All experiments were performed in biological duplicate.

### PI Assay

All measurements took place in a clear flat bottom Corning 96 well plate (cat#CLS2585 from Millipore Sigma). Assay conditions were adapted from Boix-Lemonche *et. al.^8^ B. subtilis* and *E. coli* cells were grown as described above, spun down (4,000x*g*, 10 min) and resuspended in ½ culture volume of 25 mM glucose in 1X PBS (pH = 7.4) to reach an OD_600_ ~1. A working stock of PI was made to a concentration of 1 mg/mL and diluted to a final concentration of 0.05 mg/mL into a final well volume of 100 μL. Stock solutions of the polymers were diluted to final concentrations of 0.1 mg/mL **ODB**, 0.073 mg/mL **DB**, 0.068 mg/mL **OD**, or 0.05 mg/mL **D**, 200 μM PMBN, 40 μM SNTT, 10 μg/mL colistin,^63–64^ or 2 mg/mL lysozyme. Untreated cells served as a negative control. Cells were added immediately prior to measurement at λex/λem 595/617 ± 20 nm on a Tecan Spark Plate Reader after 10 min with orbital shaking (5 ms) immediately prior to the measurement. Relative PI Uptake was calculated by the equation below where *RFU_treated_* is the measured fluorescence of the permeabilization agent treated samples, *RFU_Untreated_* is the measured fluorescence of PI in buffer, and *RFU_positive_* is the measured fluorescence of the respective positive control samples (lysozyme or colistin).

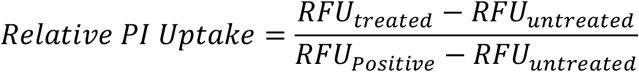

### Outer Membrane Damage Assay

Overnight cultures of *E. coli* BL21 pHisǁ cells were diluted 1:10 into fresh LB media with 100 μg/mL ampicillin and grown to OD_600_ ~0.5, cells were spun down (4,000x*g*, 10 min), supernatant removed, and cell pellets were resuspended in ½ culture volume of 5 mM HEPES buffer (pH = 7.2) to OD_600_ ~1.0. A stock solution of 0.5 mM NPN in acetone was diluted to 40 μM in a clear flat bottom Corning 96 well plate (cat#CLS2585 from Millipore Sigma) with 5 mM HEPES buffer. All other compounds were diluted to their final concentrations of 0.1 mg/mL **ODB**, 0.073 mg/mL **DB**, 0.068 mg/mL **OD**, or 0.05 mg/mL **D** polymer, 200 μM PMBN, 40 μM SNTT, and 0.01 mg/mL colistin. Cells were added immediately prior to measurement at λex/λem 355/405 ± 20 nm on a Tecan Spark Plate Reader. Measurements were taken at 10 min intervals with orbital shaking (5 ms) immediately prior to reading. Relative NPN Uptake was calculated at 10 min into measurement by the following equation:

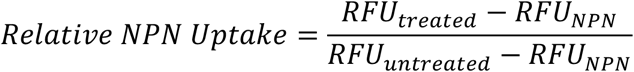

Where RFU_treated_ is the measured fluorescence of treated samples, the RFU_NPN_ is the measured fluorescence of the NPN solution alone, and RFU_untreated_ is the measured fluorescence of the bacteria with NPN dye. All measurements were taken in technical triplicate and biological duplicate.

### Resazurin Assay

A 1000X working stock of resazurin sodium salt (12.5 mg/mL) in LB media was filter-sterilized through a 0.22 μM PES filter, then diluted with LB media to a 12.5 μg/mL working stock solution. This working stock was pipetted into each well to a volume of 190 μL. Cells were grown as previously mentioned, aliquots were spun down (2,000x*g*,1 min; 500 μL of *E. coli*, 1 mL of *B. subtilis*) and supernatant removed. Each sample was resuspended in the designated solutions of 0.1 mg/mL **ODB**, 0.073 mg/mL **DB**, 0.068 mg/mL **OD**, 0.05 mg/mL **D**, 200 μM PMBN, 40 μM SNTT, 10 μg/mL colistin, or 2 mg/mL lysozyme in 30 mM TES buffer (pH = 7.4) to a total volume of 40 μL and incubated (37 °C, 30 min). Cells were centrifuged (2,000x*g*, 1 min), supernatant removed, and cells were washed once with TES buffer (100 μL). Treated cells were then resuspended in TES buffer (100 μL) and 10 μL of this sample was added into each well for a total volume of 200 μL. The 96-well plate was incubated (37 °C, 10 min) in the dark and all measurements took place in a clear flat bottom Corning 96 well plate (cat#CLS2585 from Millipore Sigma) with measured fluorescence at λex/λem 535/580 ± 20 nm on a Tecan Spark Plate Reader. Relative fluorescence intensity was calculated according to the following equation:

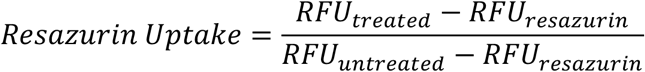

where RFU_treated_ is the measured fluorescence value of the treated samples, RFU_resazurin_ is the measured fluorescence value of resazurin solution alone without cells, and RFU_untreated_ is the measured fluorescence value of the cells in buffer. All measurements were done in technical triplicate and biological duplicate.

### Live-Cell Imaging

*E. coli* cells were grown as described in the cell culture section and treated according to the gel-based assay protocol, stopping before cell lysis. An agarose pad was made with 0.1% low melting agar in LB media, then slowly poured into the center of a slide containing an outer, rectangular border of double-sided tape. It was set aside to cool at 4 °C for at least 30 min prior to use. After treatment with polymers, the cells were resuspended in fresh TES (pH = 7.4) and pipetted through a 1 ½ in. 20-gauge needle eight times to disperse any clumps. The sample (20 μL) was transferred to the center of the agarose pad and the slide was slowly rotated to disperse cells completely across the pad. A cover slip (24×40 mm) was pressed into the sides of the double stick tape. Samples were imaged in bright field and the TRITC channel (λex/λem 545/576 ± 25 nm) with a Cell Vivo environmental chamber maintained at 37 °C and an ambient atmosphere, using an Olympus IX-73 microscope and oil immersion on a UPlanSApo 100x/1.40 Oil Ph3/0.17/FN26.5 objective lens. Micrographs were captured with a Hamamatsu Orca Flash 4.0 LT C11440 – 42U30 CMOS camera. Exposure time was ~92 ms. Growth of the cells was monitored by taking images every 15 min for at least 2 h, approximately 10x one division cycle.

