## Supporting Information Document for "Cationic Polymers Enable Internalization of Negatively Charged Chemical Probe into Bacteria"

**Experimental Methods**

General Materials and Methods

SDS-PAGE analysis

Dynamic Light Scattering

Electrophoretic Mobility Shift Assay

Bacterial Growth Assay

**Supporting Figures, Tables, and Movies**

**Figure S1:** Native *E. coli* BL21 fluorescent protein.

**Scheme S1:** Scheme for **B-TMR-ATPγS** probe synthesis.

**Figure S2:** Concentration-dependent labeling of HK853 with **B-TMR-ATPγS**.

**Figure S3:** Determination of the optimal nitrogen to phosphorous ratio for the internalization of ATP derivatives.

**Figure S4:** Comparison of *E. coli* BL21 treatment with **B-TMR-ATPγS** at different probe concentrations.

**Table S1:** HK853 labeling by **B-TMR-ATPγS** compared to coomassie staining (total protein) following treatment with **B-TMR-ATPγS** and permeabilization reagent.

**Figure S5:** *E. coli* BL21 treatment with **B-TMR-ATPγS** and permeabilization reagents.

**Figure S6:** *E. coli* BL21 treated with **O** polymer and **B-TMR-ATPγS**.

**Figure S7:** Impact of BODIPY on internalization of ATP derivatives into bacterial cells and proteome labeling in *E. coli* BL21 cells.

**Figure S8:** *E. coli* and *B. subtilis* growth with polymer treatment.

**Figure S9:** Inherent fluorescence of each polymer.

**Figure S10:** Resazurin uptake with colistin.

**Figure S11:** Live Cell Imaging, treated with **D** polymer and **B-TMR-ATPγS**.

**Figure S12:** Live Cell Imaging, treated with **B-TMR-ATPγS** only.

**Figure S13:** Live Cell Imaging, treated with **DB** polymer only.

**Figure S14:** Live Cell Imaging, treated with **D** polymer only.

**Figure S15:** Live Cell Imaging, treated with PMBN and **B-TMR-ATPγS**.

**Figure S16:** Live Cell Imaging, treated with PMBN only.

**Figure S17:** Live Cell Imaging, treated with SNTT and **B-TMR-ATPγS**.

**Figure S18:** Live Cell Imaging, treated with SNTT only.

**Table S2:** HK853 labeling for pre- and co-treatment with **B-TMR-ATPγS** and polymers compared to total protein.

**Figure S19:** Coomassie Stain for gel in Figure 4.

**Figure S20:** HK853 labeling for pre- and co-treatment with **B-TMR-ATPγS** and PMBN or SNTT.

**Figure S21:** DLS measurements of **B-TMR-ATPγS** with polymers.

**Table S3:** Normalized intensity counts for DLS measurements.

**Figure S22:** Electrophoretic mobility shift assay of **B-TMR-ATPγS** with polymers.

**Figure S23:** Comparison of ATP alone and ATP co-incubated with OD polymer by $^{31}$P NMR.

**Figure S24:** Overlay of $^{31}$P NMR spectra of ATP and ATP co-incubated with OD polymer.

**Figure S25:** Coomassie Stain for gel in Figure 5A.

**Table S4:** Summary of assay results for each permeabilization reagent.

**Figure S26:** $^{1}$H NMR of **B-TMR-ATPγS**.

**Figure S27:** $^{31}$P NMR of **B-TMR-ATPγS**.

<u>**Available as separate files:**</u>

**Movie S1:** Live Cell Imaging, treated with **DB** polymer and **B-TMR-ATPγS**. File Name: DB Polymer and Probe.mp4

**Movie S2:** Live Cell Imaging, treated with **DB** polymer only. File Name: DB Polymer Control.mp4

**Movie S3:** Live Cell Imaging, treated with **D** polymer and **B-TMR-ATPγS**. File Name: D Polymer and Probe.mp4

**Movie S4:** Live Cell Imaging, treated with **D** polymer only. File Name: D Polymer Control.mp4

**Movie S5:** Live Cell Imaging, treated with **B-TMR-ATPγS** only. File Name: Probe Control.mp4

**Movie S6:** Live Cell Imaging, treated with PMBN and **B-TMR-ATPγS**. File Name: PMBN and Probe.mp4

**Movie S7:** Live Cell Imaging, treated with PMBN only. File Name: PMBN Control.mp4

**Movie S8:** Live Cell Imaging, treated with SNTT and **B-TMR-ATPγS**. File Name: SNTT and Probe.mp4

**Movie S9:** Live Cell Imaging, treated with SNTT only. File Name: SNTT Control.mp4

*Materials.* All chemicals were obtained through the following companies. LB Media (Sigma, powder, cat #L3022), Agar (Sigma, powder, cat #A1296), APS (BioRad, 161-0700), 40% Acrylamide (BioRad, 161-0146), TEMED (BioRad, 161-0800), TrisHCl Stacking (BioRad, 1610799), TrisHCl Resolving (Biorad, 1610798), 1.5 mm Cassettes (ThermoFisher, NC2015), PageRuler Broad Range Unstained Protein Ladder (ThermoFisher, Cat # 26630), Coomassie brilliant blue R-250 staining solution (Bio-Rad, cat #1610436), Plastic 17 x100 mm Culture tubes (25 pack) (VWR, cat#60818-703), 96 well plate, clear (Millipore Sigma, cat#CLS2585).

*SDS-PAGE.* Polyacrylamide gels were made using Biorad 1.5 mm mini-protean gel cassettes and composed of a 10% resolving gel [21 mL of MilliQ, 10.5 mL of 1.5 M Tris-HCl buffer pH 8.8, 10.5 mL of acrylamide:bis-acrylamide 29:1 (40% solution, Bio-Rad), 140 µL of aqueous 10% w/v ammonium persulfate (APS), 15 µL of tetramethylethylenediamine (TEMED, Bio-Rad) added in that order], which was poured to 3/4 of the gel cassette height followed by 200 µL of ethanol and allowed to polymerize for one hr. Ethanol was removed and a 4.5% stacking gel [2.5 mL of 0.5 M Tris-HCl buffer pH 6.8, 1.125 mL of acrylamide:bis-acrylamide 29:1 (40% solution, Bio-Rad), 6.375 mL of $H_2O$, 30 µL of aqueous 10% w/v APS, 10 µL of TEMED in that order] was added, followed by a 15 well comb and the stacking portion was polymerized for 3 hrs. Amounts listed here make four gels. Gels were chilled (4 °C) during electrophoresis with the following parameters: 180 V, 40 mA and 60 W in 1X Tris running buffer (diluted from 10X stock) for 1 hr.

*SDS-PAGE loading buffer.* 4X SDS-PAGE sample loading buffer containing 200 mM Tris-HCl (pH=6.8), 40% glycerol, 8% SDS (w/v), 4% β-mercaptoethanol, and 0.8% bromophenol blue (w/v) was used.

*In-gel fluorescence detection.* After SDS-PAGE, gels were washed three times with MilliQ water, and scanned on a GE Typhoon FLA 9500 gel scanner in the TAMRA channel (λex/λem 542 nm/568 nm) or BODIPY channel (λex/λem 504 nm/514 nm).

*Coomassie Staining.* Immediately following fluorescence detection, gels were microwaved for 1 min in Coomassie Brilliant blue R-250 Staining Solution (cat# 1610436) and incubated in this solution for 30 min. Gels were then removed from Coomassie solution, washed three times with MilliQ water and destained overnight with an aqueous solution of 10% acetic acid and 40% methanol on an orbital rocker.

If further destaining was required, destaining solution was removed, replaced with MilliQ water and left for an additional 24 hrs on an orbital rocker. Gels were then scanned with the same Typhoon gel scanner in the Coomassie channel (λex/λem 532 nm/570 nm).

*Dynamic Light Scattering (DLS).* DLS solutions were analyzed on a Wyatt DynaPro III instrument (Wyatt Technology, Santa Barbara, CA). Each well was scanned 8 times with 8 s per acquisition.

*Electrophoretic Mobility Shift Assay.* Samples incubated for 30 min at RT prior to dilution with 1.33 µL of 6X loading buffer and 5 µL loaded into each well and run on a 1% agarose gel for 20 min at 150 V.

*Bacterial Growth Assay.* Overexpression strain of *E. coli* BL21 containing the pHis‖ HK853 plasmid was streaked onto LB agar plates containing 100 µg/mL ampicillin from a frozen glycerol stock and grown overnight at 37 °C. From these plates, a single colony was inoculated into fresh Lennox Broth (LB) media containing 100 µg/mL ampicillin and grown overnight at 37 °C, shaking at 220 RPM. A 1:100 dilution of the overnight culture was performed in fresh LB media containing 100 µg/mL ampicillin and grown until $OD_{600}$ ~0.7 (Genesys 30 Visible Spectrophotometer). A one mL aliquot was taken from the secondary culture and was serially diluted 1:100 in fresh LB media for a complete range of aliquots from 1x$10^{-1}$-1x$10^{-6}$. Each dilution was then spotted (100 μL) onto LB agar plates containing 100 μg/mL ampicillin as a control in addition to plates containing the **D** and **OD** polymer for a final concentration of 0.068 mg/mL and 0.05 mg/mL, respectively. Plates were grown overnight at 37 °C. Colony counting to obtain CFU was not possible as polymers caused beading of the media upon addition of the initial bacterial dilutions and "smearing" was evident upon growth of the bacteria. However, it was evident that bacteria were able to grow similarly between the polymer-treated and control conditions. Identical steps were taken with *B. subtilis* 3610 with LB media. All samples run in biological triplicate and technical duplicate.

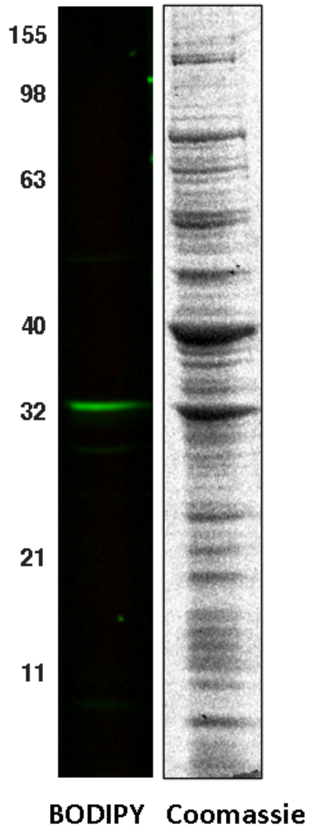


**Figure S1: Native *E. coli* BL21 fluorescent protein.** A band present in the BODIPY channel in the TES buffer control is a natively-fluorescent protein. MW Ladder in kDa.

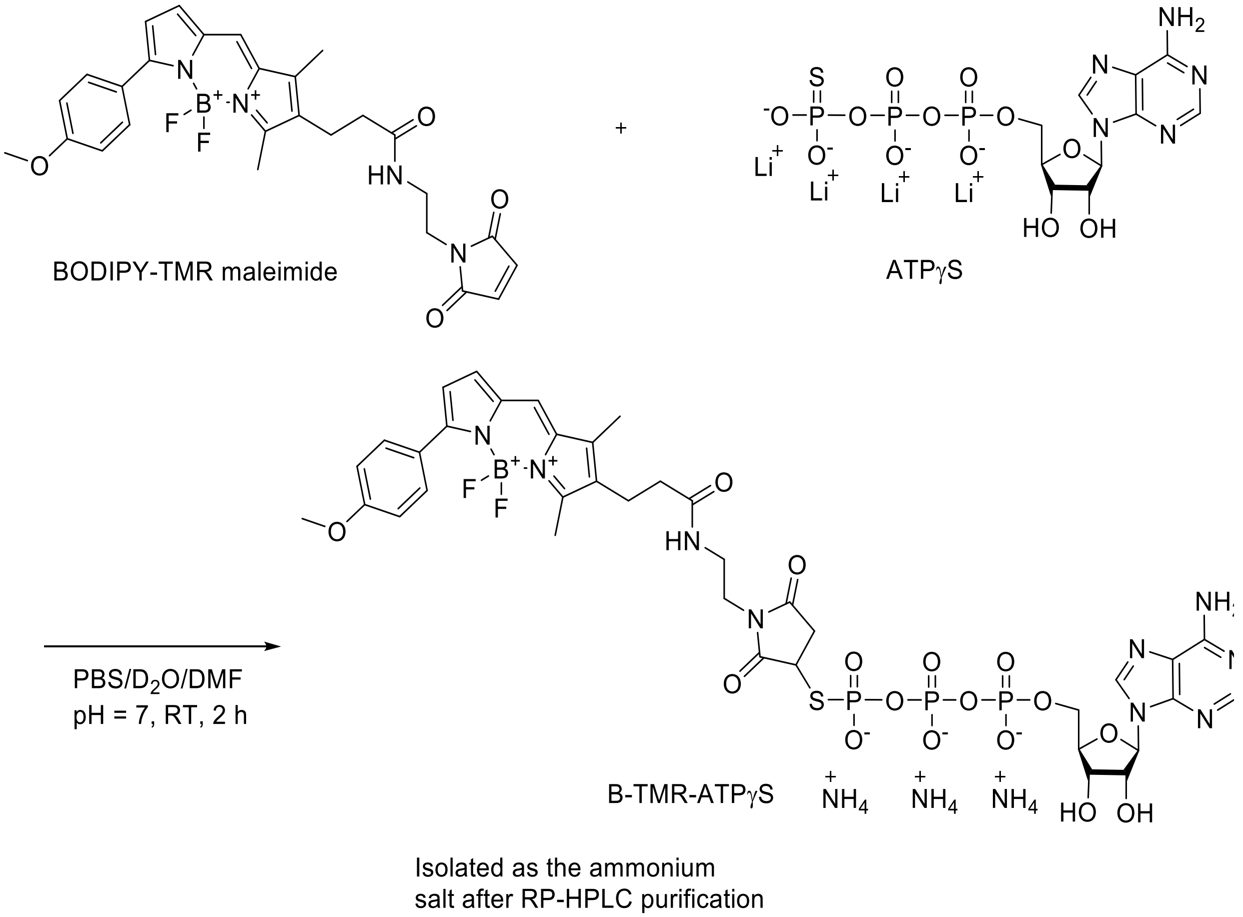


**Scheme S1: Synthesis of B-TMR-ATPγS**

*Synthesis of **B-TMR-ATPγS***. ATPγS lithium salt (2 mg, 364 µL, 3.66 mmol) was dissolved in a mixture of $D_2O$ (1.4 mL), PBS (75 mM, 186 µL), and DMF (300 µL) followed by the addition of B-TMR-maleimide in DMF (2.29 mg, 4.39 mmol). The reaction mixture was stirred at RT in the dark and deemed complete after 2 h by $^{31}P$ NMR. The reaction mixture was lyophilized and purified by RP-HPLC using an Agilent 1200 series equipped with Agilent Zorbax C3, 5 µm, 9.4 x 250 mm and a gradient of 0.1 % ammonium acetate in water (Buffer A) and 0.1% ammonium acetate in acetonitrile (Buffer B) (gradient 0-3 min 95% Buffer A; 3-12 min 95 to 5% Buffer A; 12-15 min 5% Buffer A; 3 mL/min) to afford the product **B-TMR-ATPγS**. The collected fractions were lyophilized from water several times to afford the product as the ammonium salt (88%, 3.5

mg). NMR spectra collected on a Bruker Avance III HD 400 with 5 mm BBO SmartProbe. $^{1}H$ NMR ($D_2O$, 400 MHz) δ 8.33 (d, $J$ = 2.8 Hz, 1H, d), 7.96 (s, 1H, m), 7.74 (d, $J$ = 8.4 Hz, 2H, a), 7.29 (s, 1H, n), 7.08 (d, $J$ = 6.0 Hz, 2H, b), 7.06 (d, $J$ = 4.0 Hz, 1H, d), 6.57 (d, $J$ = 4.0 Hz, 1H, l), 5.89 (m, 1H, e), 4.55 (t, $J$ = 4.8 Hz, 1H, i), 4.90 (t, $J$ = 4.4 Hz, 1H, j), 4.31 (bs, 2H, h), 4.26 (bs, 2H, h), 3.91 (s, 3H, O-$CH_3$), 3.60 (m, 2H, g), 3.36 (m, 2 H, g), 2.62 (t, $J$ = 8.0 Hz, 2H, f), 2.38 (s, 3H, $CH_3$), 2.29 (t, $J$ = 8.0 Hz, 2H, f), 2.14 (t, 3H, $CH_3$); $^{31}P$ NMR ($D_2O$, 162 MHz) 4.13 (m, 1P, γ), -11.54 (d, 1P, $J_{P\alpha,P\beta}$ = 20 Hz, α), -22.73 (m, 1P, β). δ ESI-MS (positive): $C_{37}N_9P_3O_{16}SH_{40}BF_2$ [M+H-F] = 1025.1906 (calculated), 1025.2020 (found).

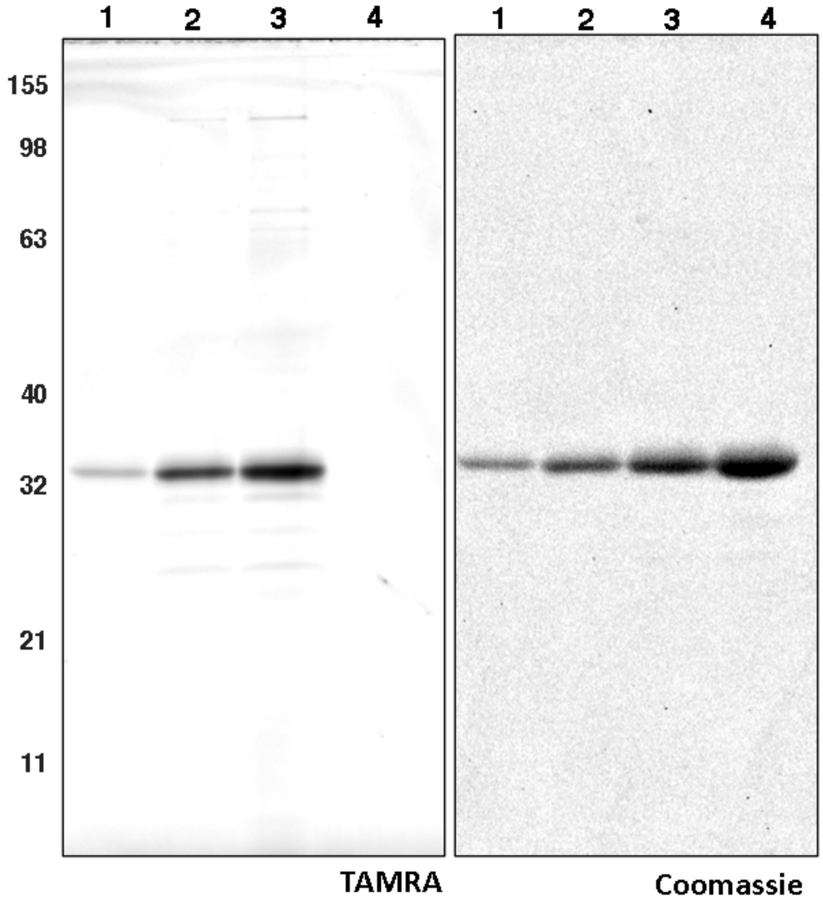


**Figure S2: Concentration-dependent labeling of HK853 with B-TMR-ATPγS.** Each lane was incubated with 1 or 2 µM HK853 (lane 1, and lanes 2 and 3, respectively) and 4, 8 or 16 µM **B-TMR-ATPγS** (lanes 1, 2 and 3, respectively) to discern if HK853 was labeled by **B-TMR-ATPγS**. Lane 4 is a protein-only control containing 4 µM HK853. Gel samples were quenched with loading buffer and then run on a 10% SDS-PAGE gel before fluorescent gel analysis. MW ladder in kDa.

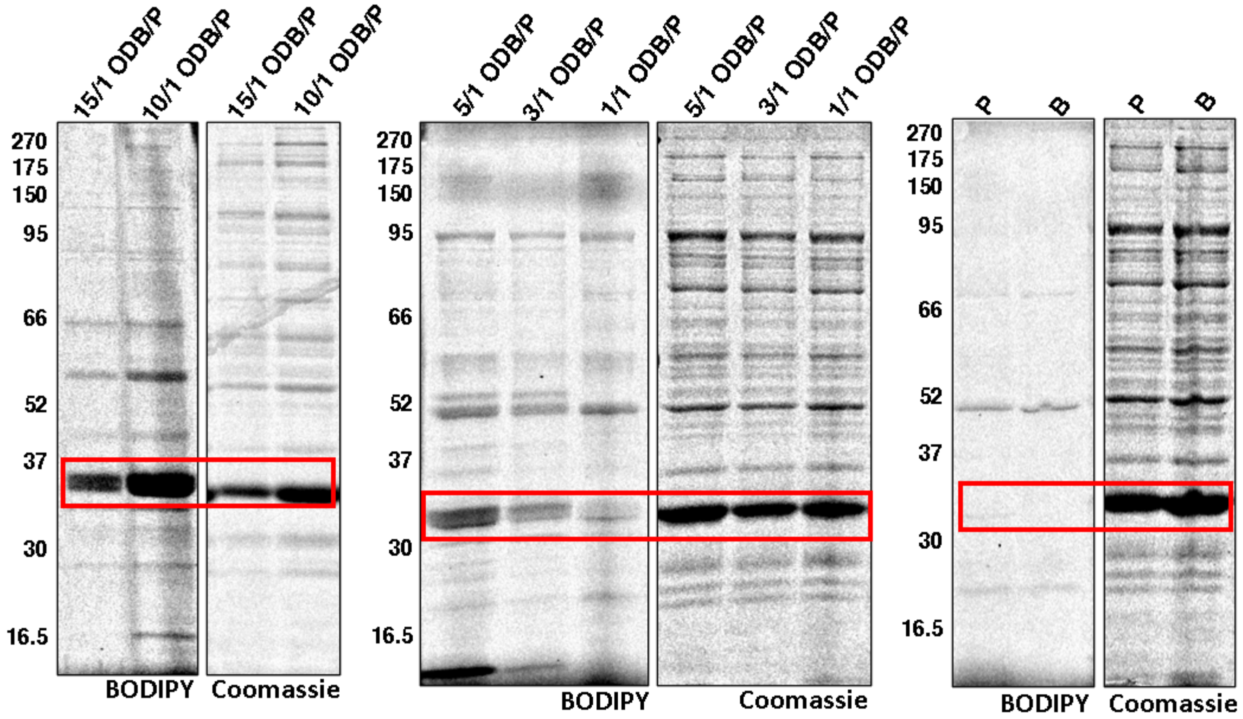


**Figure S3: Determination of the optimal nitrogen to phosphorous ratio for the internalization of ATP derivatives.** B-FL-ATPγS (P) (33 μM) was complexed with **ODB** at different N/P ratios calculated from 33 μM probe. The 1/1 N/P (0.034 mg/mL **ODB**), 3/1 N/P (0.102mg/mL **ODB**), 5/1 N/P (0.17 mg/mL **ODB**), 10/1 N/P (0.34 mg/mL **ODB**), and 15/1 N/P (0.68 mg/mL **ODB**) were mixed in water and incubated for 30 min prior to *E. coli* treatment for 1 h at 37 °C. Controls with only **B-FL-ATPγS** (P) or water (B) were also performed. After treatment, cells were spun down (2000 x *g*, 1 min) and the supernatant discarded. Cells were washed with PBS (2 x 100 μL), lysed in PBS 1% SDS, and the lysates were run on a 10% SDS-PAGE gel before fluorescent gel analysis. MW ladder in kDa. Red box indicates HK853.

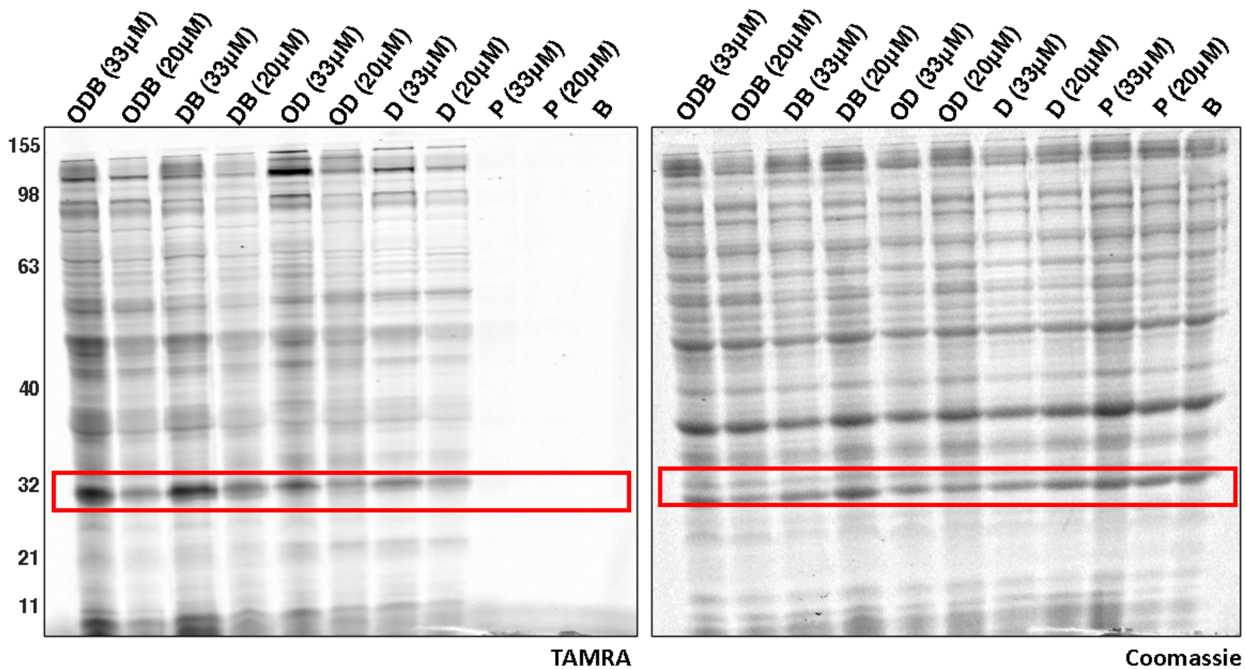


**Figure S4: Comparison of *E. coli* BL21 treatment with B-TMR-ATPγS at different probe concentrations.** Each polymer with either 33 µM or 20 µM of the probe was added to the cells at a 5/1 N/P ratio after a 30-min co-incubation. Each lane is labeled with the corresponding reagent utilized for permeabilization and either the 33 µM or 20 µM probe amount. B indicates TES buffer control. Representative gel of two biological replicates. MW ladder in kDa. Red box indicates HK853.

**Table S1: HK853 labeling by B-TMR-ATPγS compared to coomassie staining (total protein) following treatment with B-TMR-ATPγS and permeabilization reagent.** Experiment performed in technical triplicate and biological duplicate.

| Sample | Ratio ± Std Dev | Percent Labeling ± Std Dev |
|---|---|---|
| ODB | 1.01±0.07 | 100% ± 7% |
| DB | 1.01±0.09 | 100% ± 9% |
| OD | 0.94±0.03 | 94% ± 3% |
| D | 0.72±0.02 | 72% ± 2% |
| PMBN | 0.22±0.01 | 22% ± 1% |
| SNTT | 0.37±0.02 | 37% ± 2% |
| P | 0.11±0.01 | 11% ± 1% |
| B | 0.04±0.004 | 4% ± 0.4% |

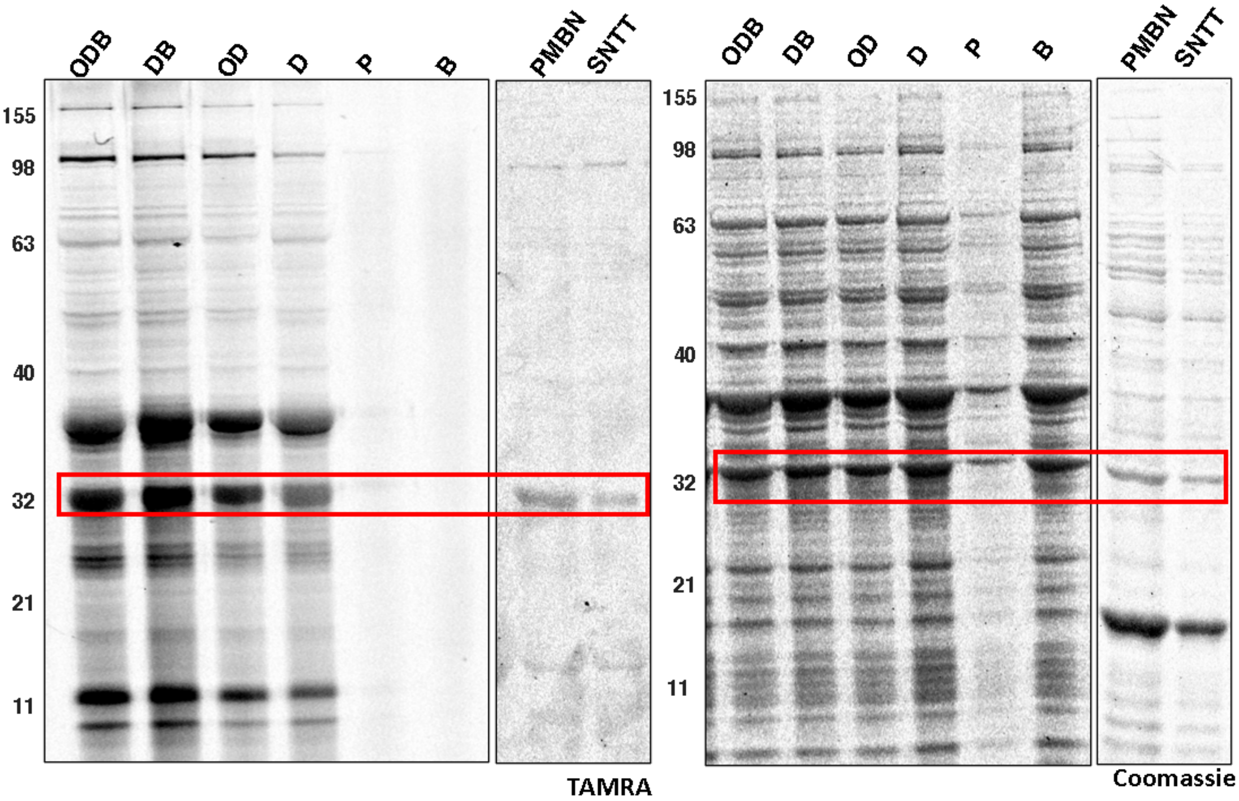


**Figure S5:** ***E. coli*** **BL21 treatment with B-TMR-ATPγS and permeabilization reagents.** Each polymer or commercially available permeabilization reagent (PMBN or SNTT) was used with 20 µM of the respective probe concentration at a 5/1 N/P ratio or 200 µM and 40 µM for PMBN and SNTT, respectively, after a 30-min co-incubation. Each lane is labeled with the corresponding reagent utilized for permeabilization. P indicates probe alone and B is TES buffer. Representative gel of biological duplicates. MW ladder in kDa. Red box indicates HK853.

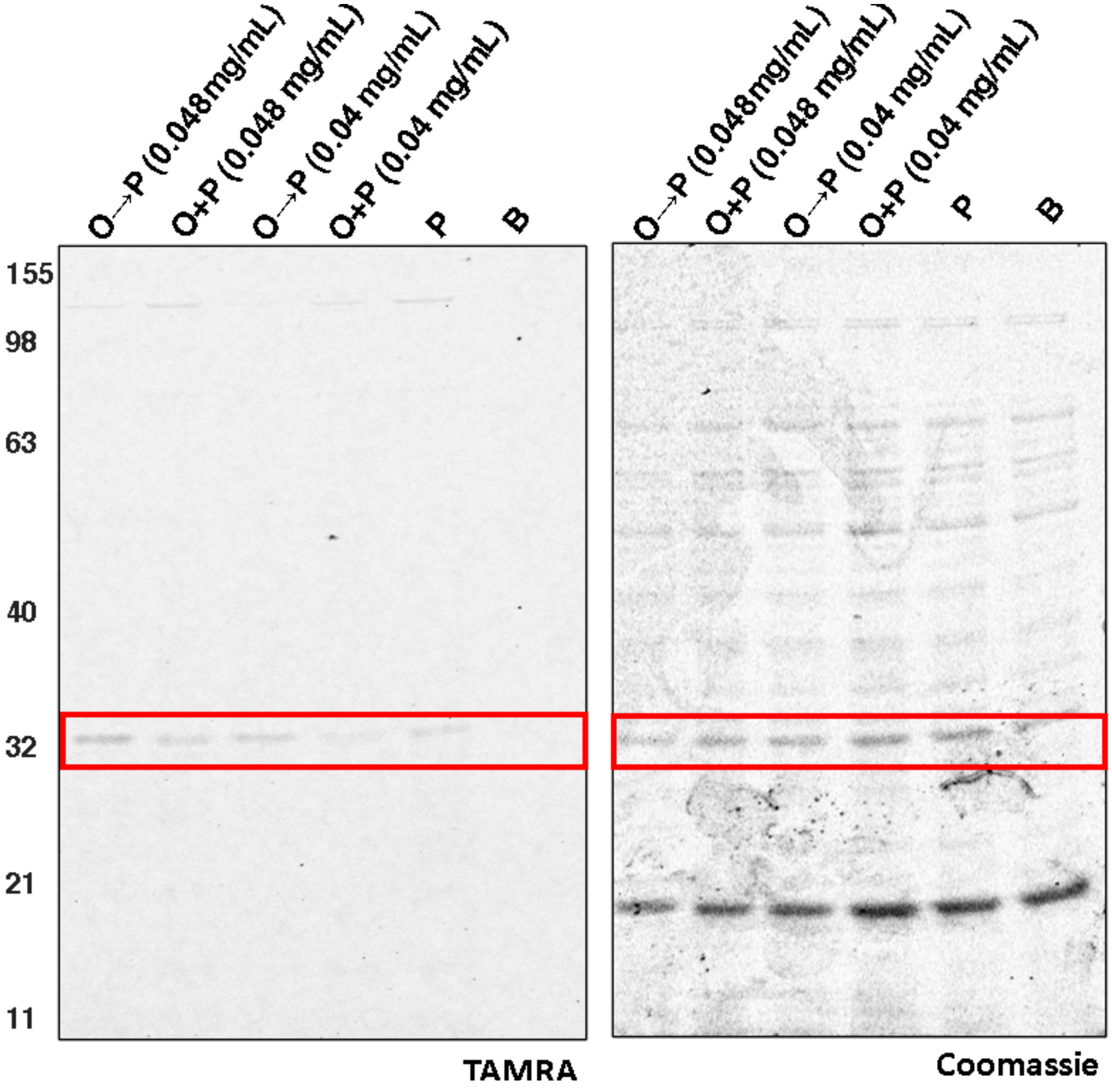


**Figure S6: *E. coli* BL21 treated with O polymer and B-TMR-ATPγS.** PEG polymer was incubated with **B-TMR-ATPγS** (20 μM) at 5/1 N/P ratios calculated for the percent of PEG found in either **ODB** (0.048 mg/mL) or **OD** (0.04 mg/mL) after a 30-min co-incubation. P stands for 20 μM probe alone and B stands for the TES buffer control. Representative gel of biological duplicates. MW ladder in kDa. Red box indicates HK853.

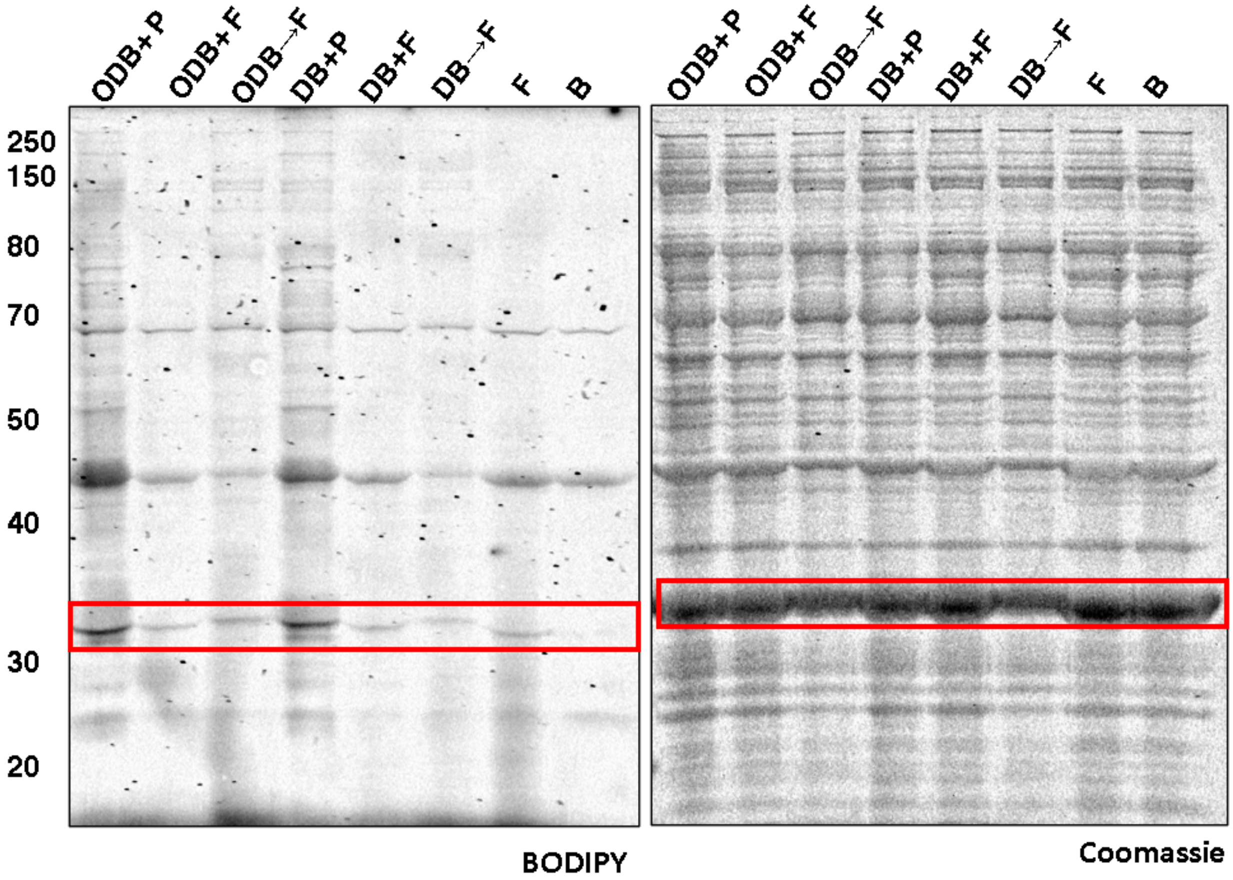


**Figure S7: Impact of BODIPY on internalization of ATP derivatives into bacterial cells and proteome labeling in *E. coli* BL21 cells.** B-FL-ATPγS (P) or BODIPY (F) (33 μM) were incubated with **ODB** (0.17 mg/mL) or **DB** (0.12 mg/mL) in water for 30 min prior to *E. coli* BL21 treatment for 30 min at 37 °C. Negative control treatments with only BODIPY (33 μM) and a water control (B) were added. Cells were also pretreated (indicated on gel with →) with **ODB** (0.17 mg/mL) or **DB** (0.12 mg/mL) for 15 min at 37 °C, washed with PBS (2 x 100 μL), and, then with BODIPY (33 μM) for 15 min at 37 °C. After treatment, cells were spun down (2000 x *g*, 1 min) and the supernatant discarded. Cells were washed with PBS (2 x 100 μL), lysed in PBS 1% SDS, and the lysates run on a 10% SDS-PAGE before fluorescence analysis. MW ladder in kDa. Red box indicates HK853.

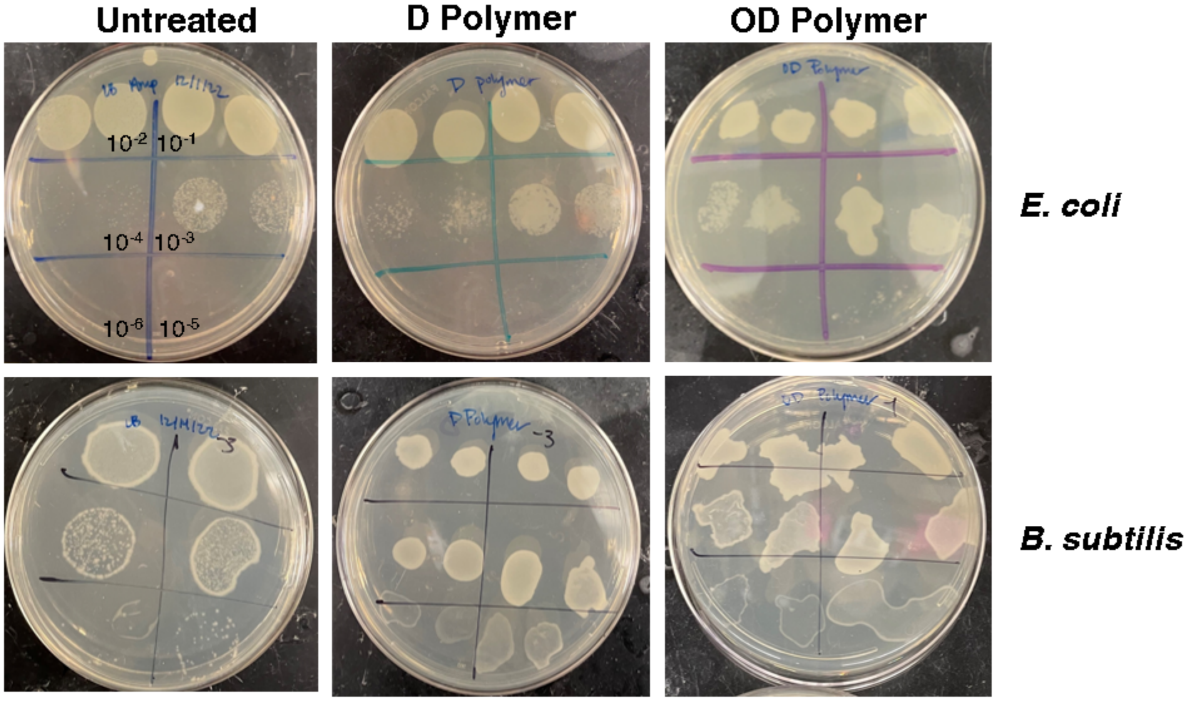


**Figure S8: *E. coli* and *B. subtilis* growth with polymer treatment.** Colony growth was measured by eye after overnight incubation as samples “smeared” due to media beading on the polymer-containing plates. Overnight cultures of *E. coli* were diluted 1:100 into fresh media and serially diluted 1:100 six times and each dilution plated onto plates containing a final concentration of 0.068 mg/mL **OD**, and 0.05 mg/mL **D**. All dilutions are labeled on the first panel for untreated *E. coli* and remain the same for each following plates. All samples were run in biological triplicate and technical duplicate (except for control *B. subtilis* for which the spots were too large).

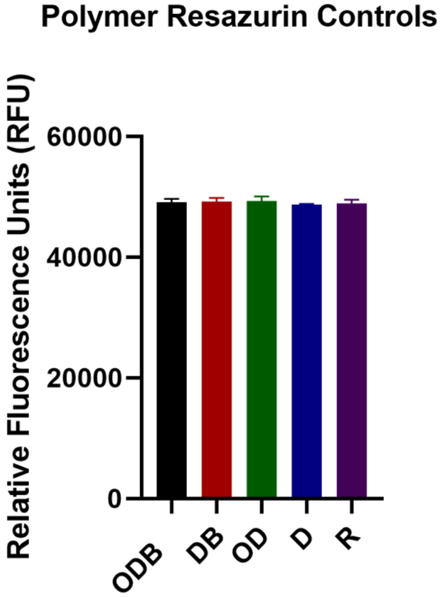


**Figure S9: Inherent fluorescence of each polymer.** Each polymer was added to give a final concentration of 0.10 mg/mL **ODB,** 0.078 mg/mL **DB**, 0.068 mg/mL **OD**, and 0.05mg/mL **D** with 190 µL of working stock resazurin (12 µg/mL) and compared to a resazurin control with a one-way ANOVA using Graphpad Prism 8, no statistically significant difference was found between each sample and the buffer control.

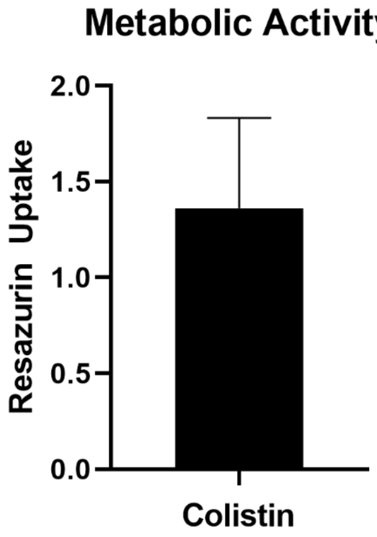


**Figure S10: Resazurin uptake with colistin.** Colistin was added to *E. coli* (BL21) for a final concentration of 0.01 mg/mL with 190 µL of working stock resazurin (12 µg/mL) and compared to a buffer-treated control. Performed in technical triplicate and biological duplicate.

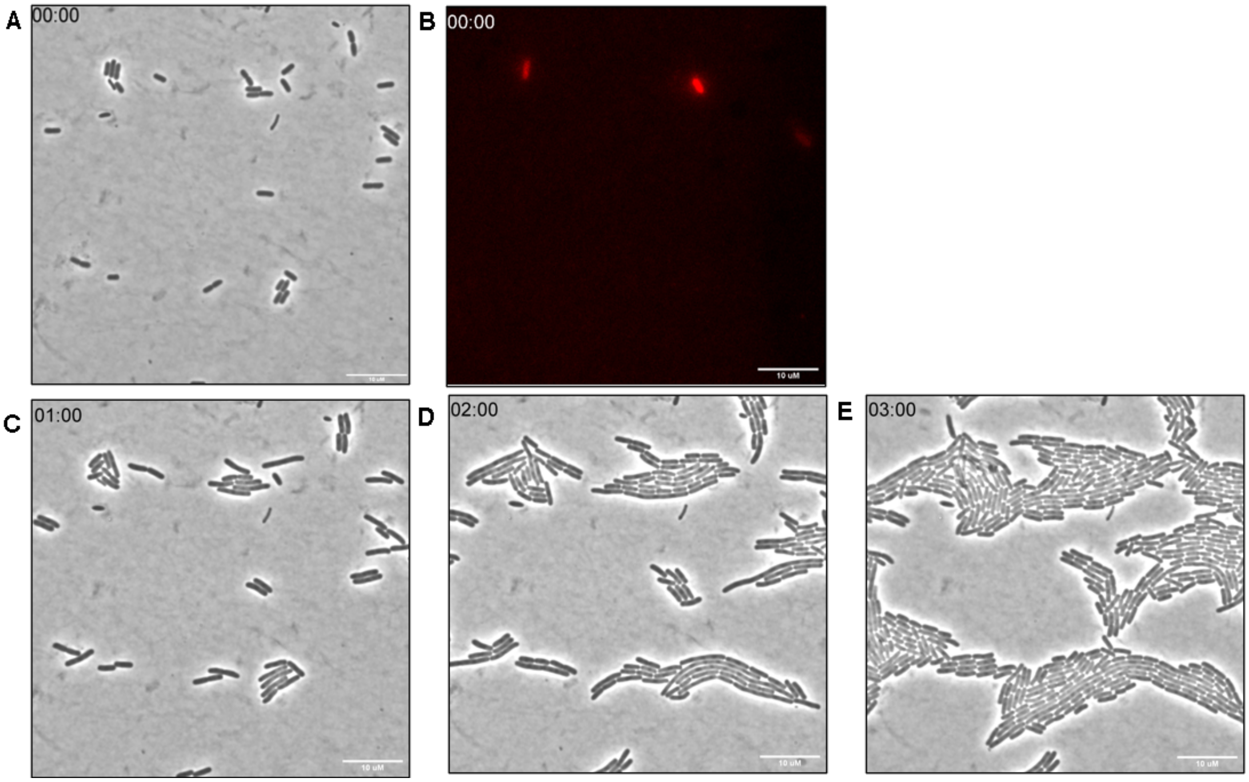


**Figure S11: Live cell imaging of *E. coli* treated with D polymer and B-TMR-ATPγS**. A. Time 0 min brightfield image. B. Time 0 min TRITC channel. C. Time 60 min brightfield. D. Time 120 min brightfield. E. Time 180 min brightfield. Scale bars are 10 μm. Movie available as separate file.

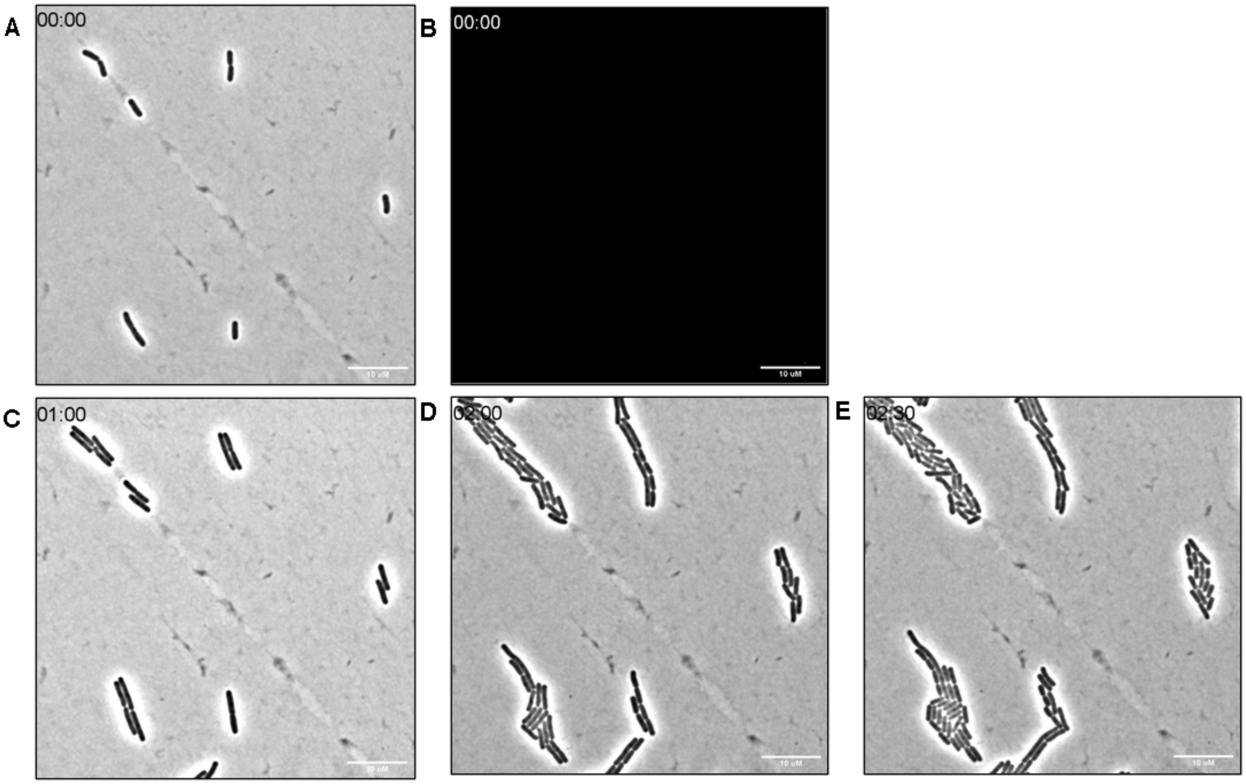


**Figure S12: Live cell imaging of *E. coli* treated with B-TMR-ATPγS only.** A. Time 0 min brightfield image. B. Time 0 min TRITC channel. C. Time 60 min brightfield. D. Time 120 min brightfield. E. Time 150 minutes brightfield. Scale bars are 10 μm. Movie available as separate file.

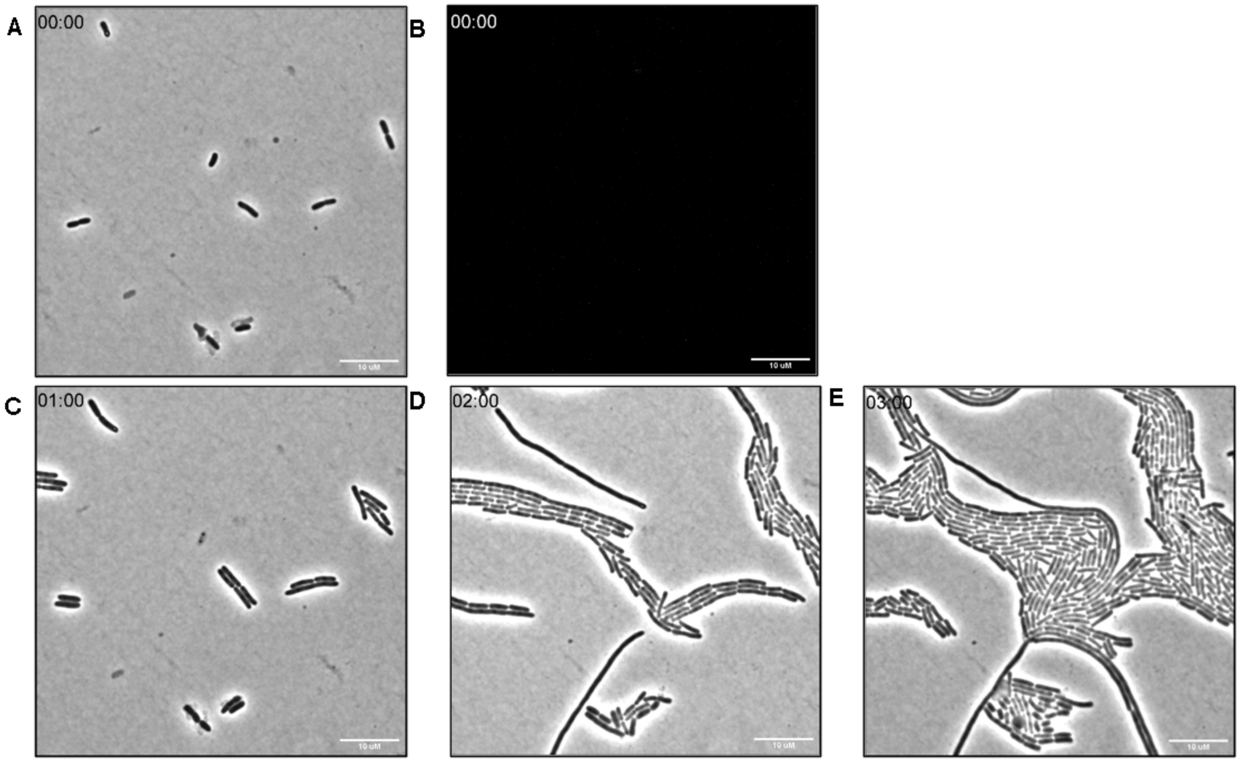


**Figure S13: Live cell imaging of *E. coli* treated with DB polymer only.** A. Time 0 min brightfield image. B. Time 0 min TRITC channel. C. Time 60 min brightfield. D. Time 120 min brightfield. E. Time 180 min brightfield. Scale bars are 10 μm. Movie available as separate file.

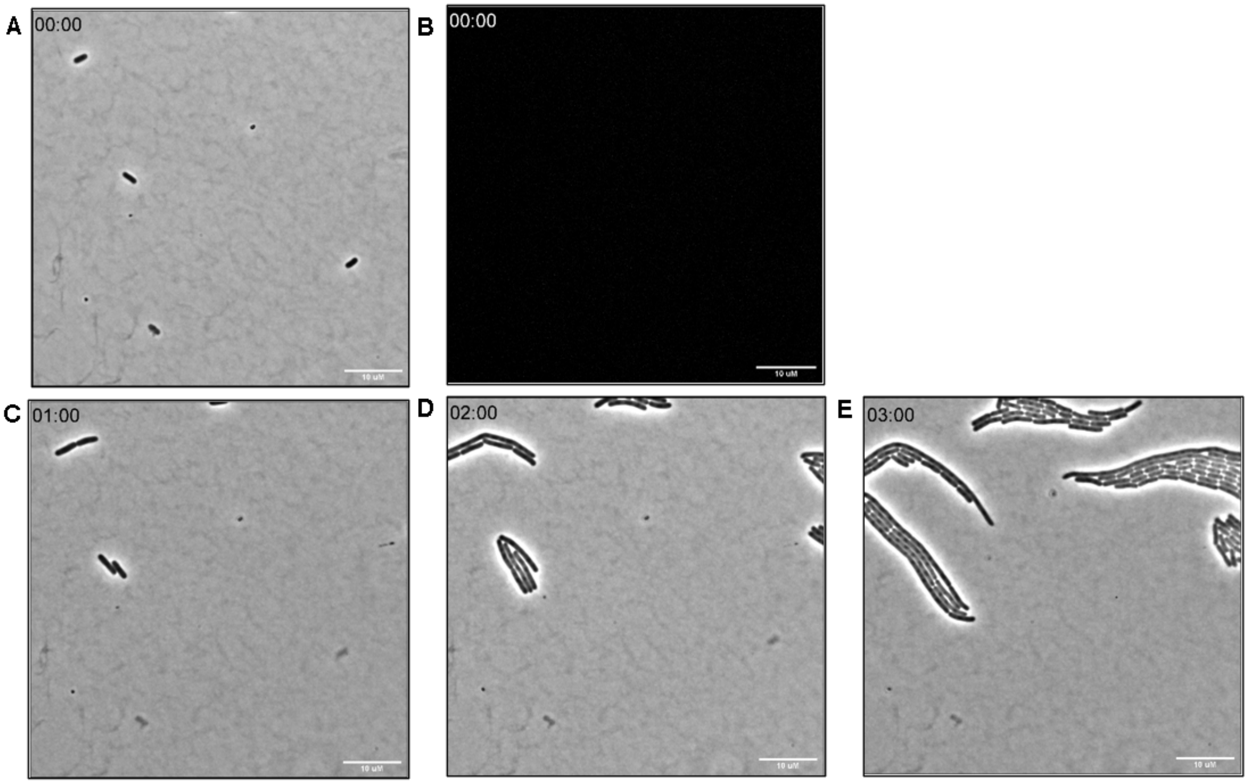


**Figure S14: Live cell imaging of *E. coli* treated with D polymer only.** A. Time 0 min brightfield image. B. Time 0 min TRITC channel. C. Time 60 min brightfield. D. Time 120 min brightfield. E. Time 180 min brightfield. Scale bars are 10 μm. Movie available as separate file.

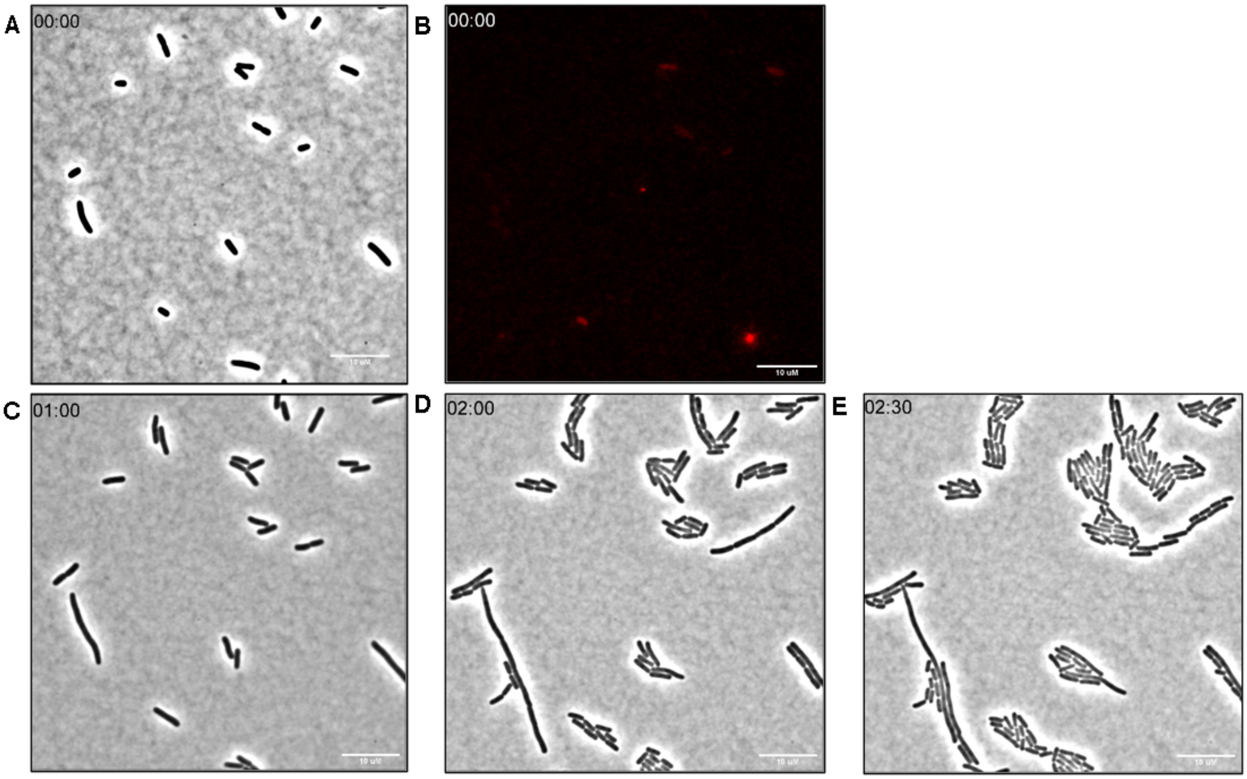


**Figure S15: Live cell imaging of *E. coli* treated with PMBN and B-TMR-ATPγS.** A. Time 0 min brightfield image. B. Time 0 min TRITC channel. C. Time 60 min brightfield. D. Time 120 min brightfield. E. Time 150 min brightfield. Scale bars are 10 μm. Movie available as separate file.

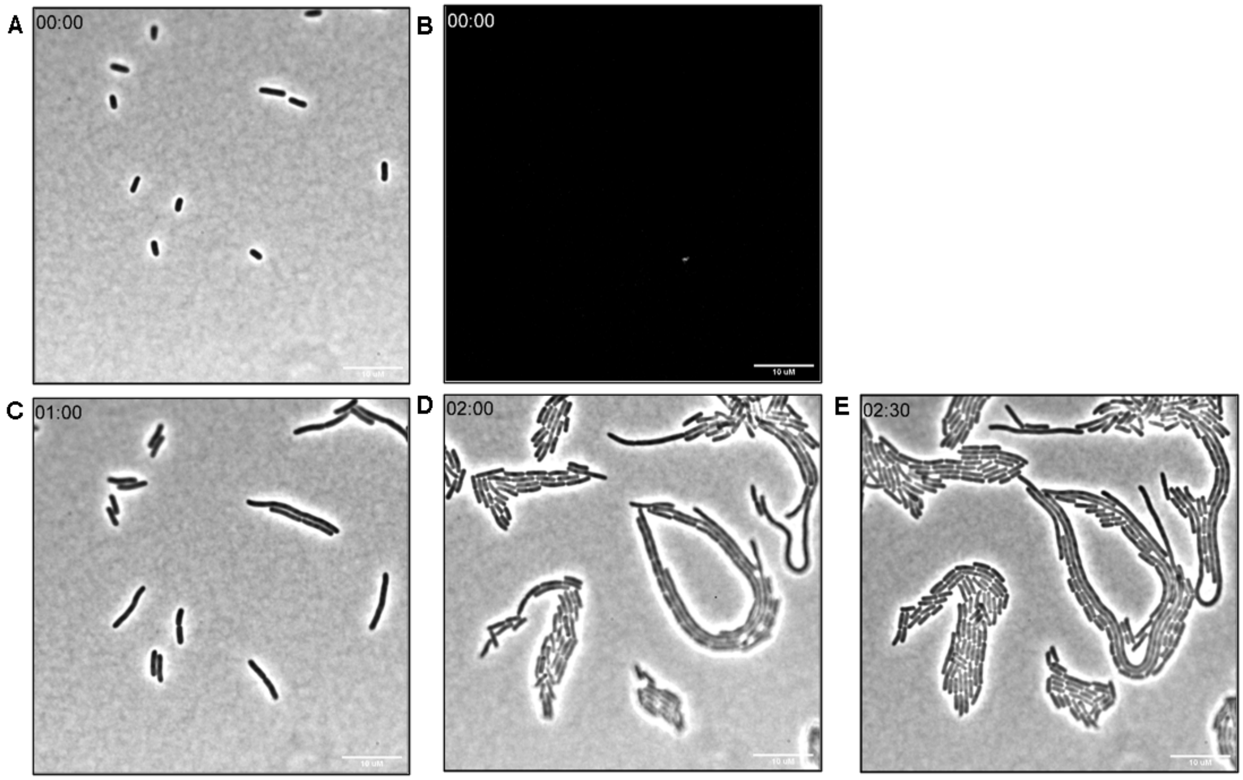


**Figure S16: Live cell imaging of *E. coli* treated with PMBN only.** A. Time 0 min brightfield image. B. Time 0 min TRITC channel. C. Time 60 min brightfield. D. Time 120 min brightfield. E. Time 150 min brightfield. Scale bars are 10 µm. Movie available as separate file.

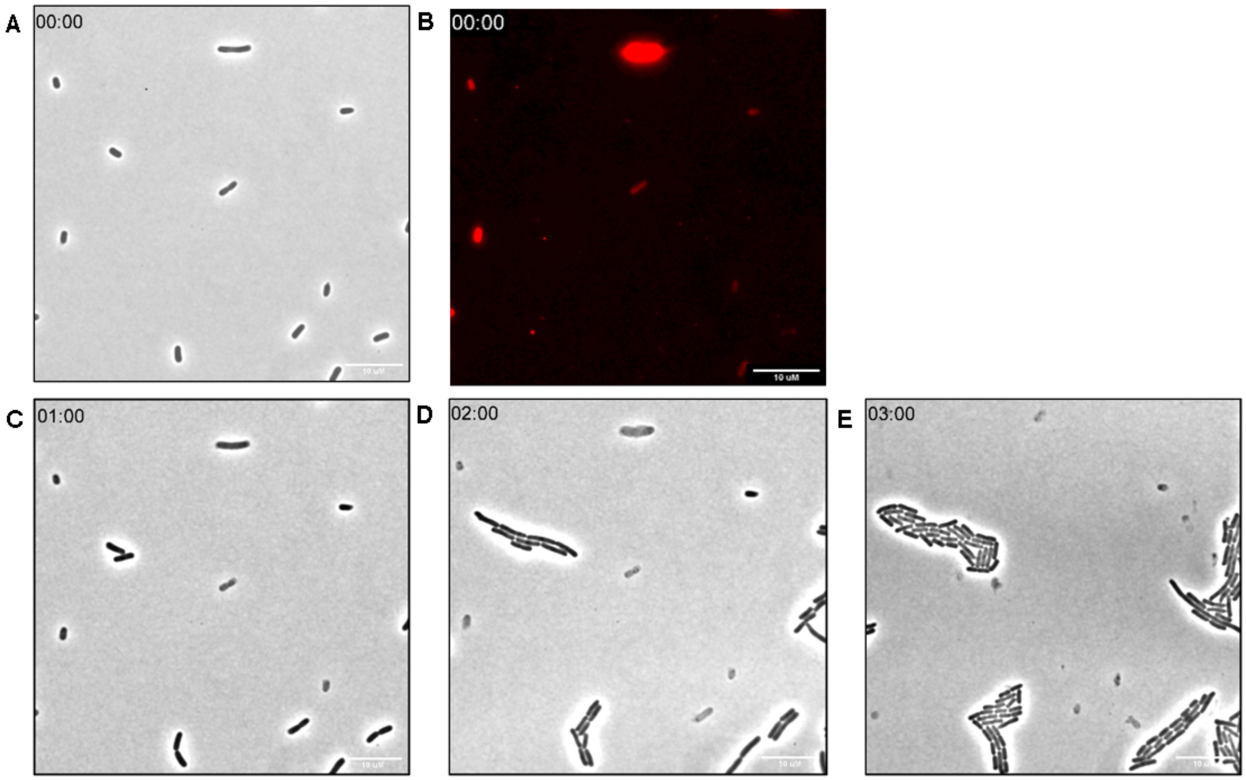


**Figure S17: Live cell imaging of *E. coli* treated with SNTT and B-TMR-ATPγS.** A. Time 0 min brightfield image. B. Time 0 min TRITC channel. C. Time 60 min brightfield. D. Time 120 min brightfield. E. Time 180 min brightfield. Scale bars are 10 μm. Movie available as separate file.

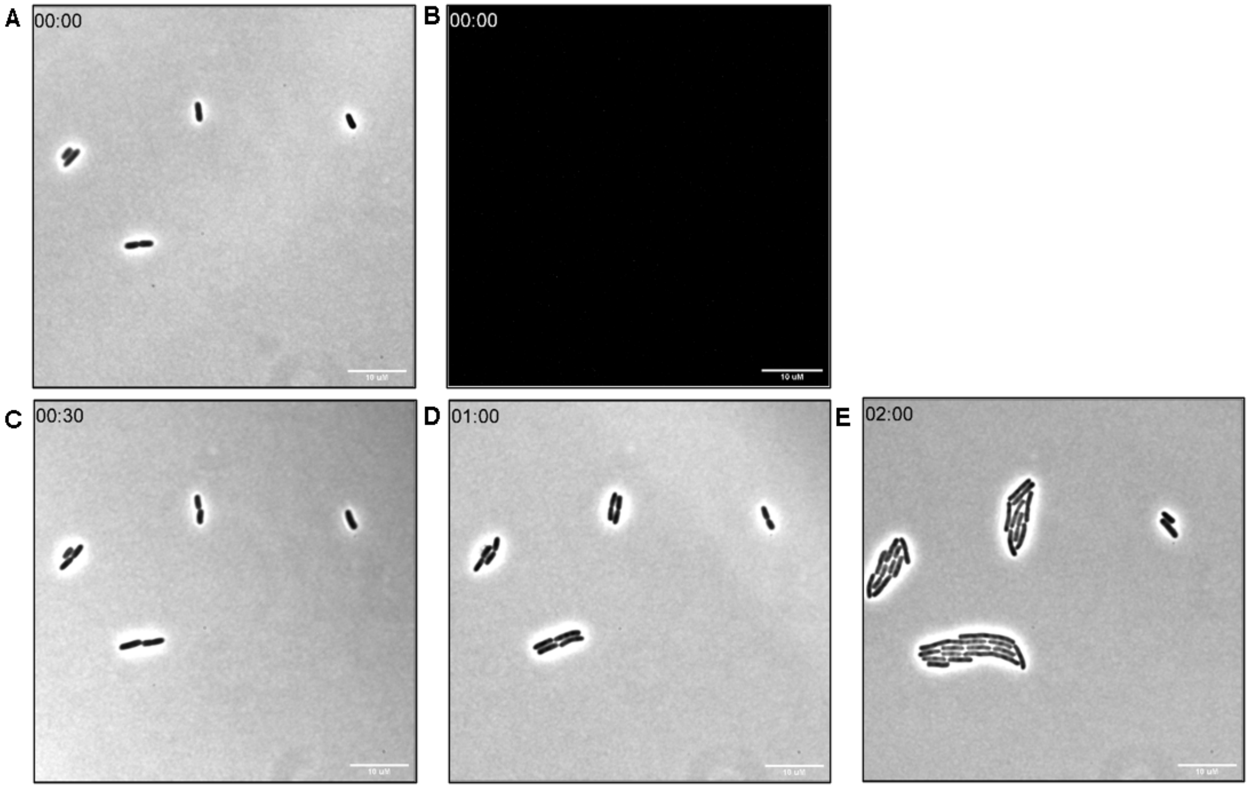


**Figure S18: Live cell imaging of *E. coli* treated with SNTT only.** A. Time 0 min brightfield image. B. Time 0 min TRITC channel. C. Time 30 min brightfield. D. Time 60 min brightfield. E. Time 120 min brightfield. Scale bars are 10 μm. Movie available as separate file.

**Table S2: HK853 labeling (fluorescence) for pre- and co-treatment of polymers with B-TMR ATPγS compared to total HK853 protein concentration (Coomassie).** Samples labeled as "pre" indicate pretreatment of polymers while "co" labeled samples indicate cotreatment of polymers. P is for the probe treatment alone, and B is the TES buffer control. Calculations performed in technical triplicate and biological duplicate.

| Sample | Ratio ± Std Dev |
| --- | --- |
| **ODB** Pre | 0.95 ± 0.02 |
| **ODB** Co | 1.34 ± 0.01 |
| **DB** Pre | 0.60 ± 0.01 |
| **DB** Co | 1.35 ± 0.02 |
| **OD** Pre | 0.74 ± 0.003 |
| **OD** Co | 1.75 ± 0.04 |
| **D** Pre | 0.94 ± 0.01 |
| **D** Co | 2.07 ± 0.01 |
| P | 0.61 ± 0.01 |
| B | 0.23 ± 0.002 |

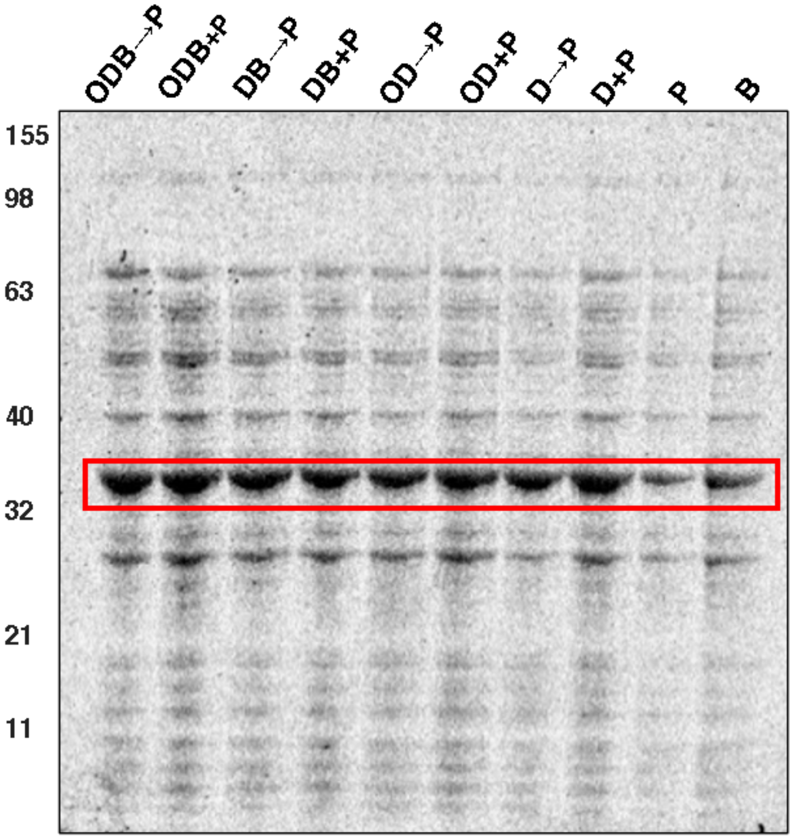


**Figure S19: Coomassie Stain of gel in Figure 4**. MW Ladder in kDa. Red box indicates HK853.

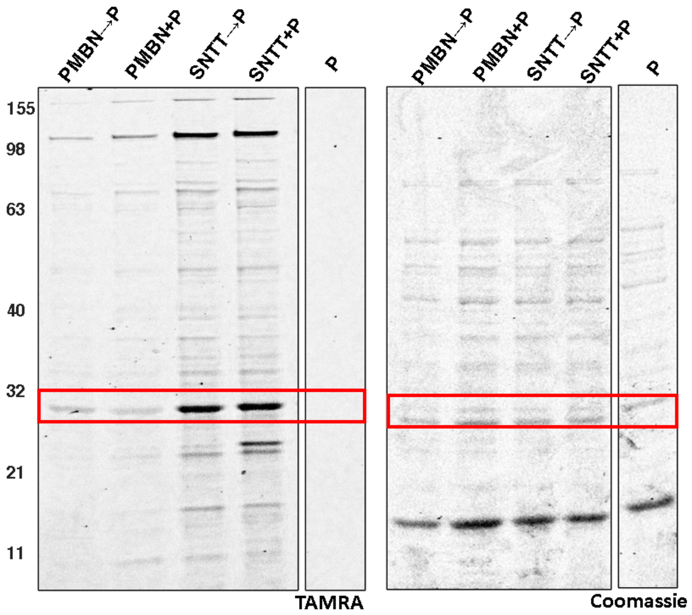


**Figure S20: HK853 labeling for pre- and co-treatment with B-TMR-ATPγS and PMBN or SNTT.** *E. coli* were pre-incubated with SNTT (40 µM) or PMBN (200µM) for 30 min and then treated with **B-TMR-ATPγS** (20 µM) for an additional 30 min. Alternatively, cells were co-treated with SNTT and PMBN with **B-TMR-ATPγS** for 30-min. P stands for 20 µM probe alone and B stands for the TES buffer control. Representative gel of biological triplicates. MW ladder in kDa. Red box indicates HK853.

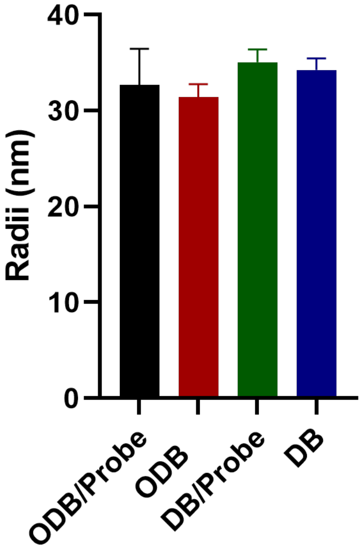


**Figure S21: DLS measurements of B-TMR-ATPγS with polymers.** Solutions were prepared in sterile water at the optimized 5/1 N/P ratio for each polymer (**ODB, DB, OD,** or **D**) to 40 μM probe (Probe). Samples mixed at RT for 30 min prior to analysis. Statistical differences in hydrodynamic radii were evaluated by one-way ANOVA in Graphpad Prism 8.0.

**Tables S3: Normalized intensity counts for DLS measurements.** Averaged kilacounts (kCnt) for **OD** and **D** polymer samples measured with and without **B-TMR-ATPγS**.

| Normalized Intensities (kCnt/s) | | |
|---|---|---|
| **Sample** | **With Probe** | **Without Probe** |
| **OD** | 7798 ± 2990 | 4841 ± 2494 |
| **D** | 10552 ± 3724 | 4315 ± 2676 |

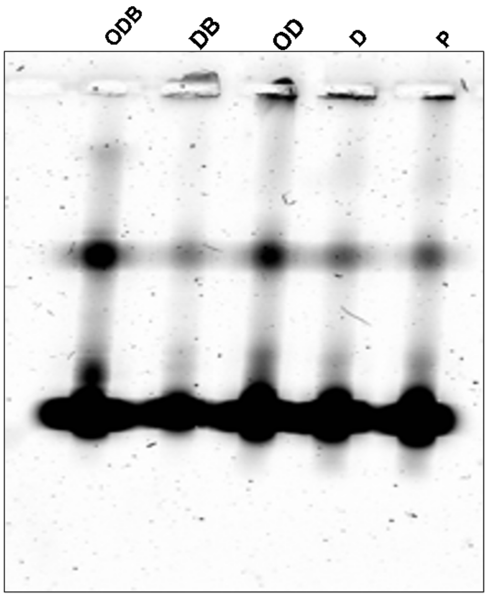


**Figure S22: Electrophoretic mobility shift assay.** Each polymer (**ODB, DB, OD,** or **D**) was incubated with **B-TMR-ATPγS** probe (P) at a 5/1 N/P ratio for a final concentration of **B-TMR-ATPγS** probe at 106 μM, 10 μL of sample was loaded into the gel for analysis.

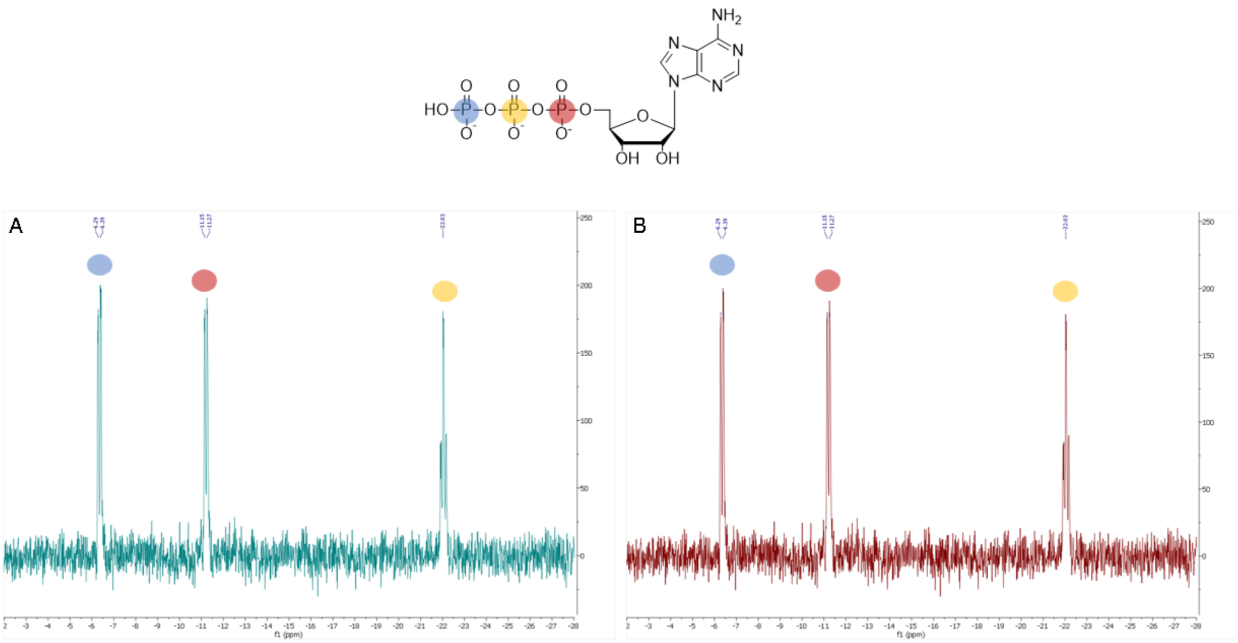


**Figure S23: Comparison of ATP alone and ATP co-incubated with OD polymer by $^{31}$P NMR.** A. $^{31}$P NMR spectra of 200 μM ATP alone. B. $^{31}$P NMR spectra of 200 μM ATP incubated with 0.4 mg/mL OD polymer for 30 min at RT.

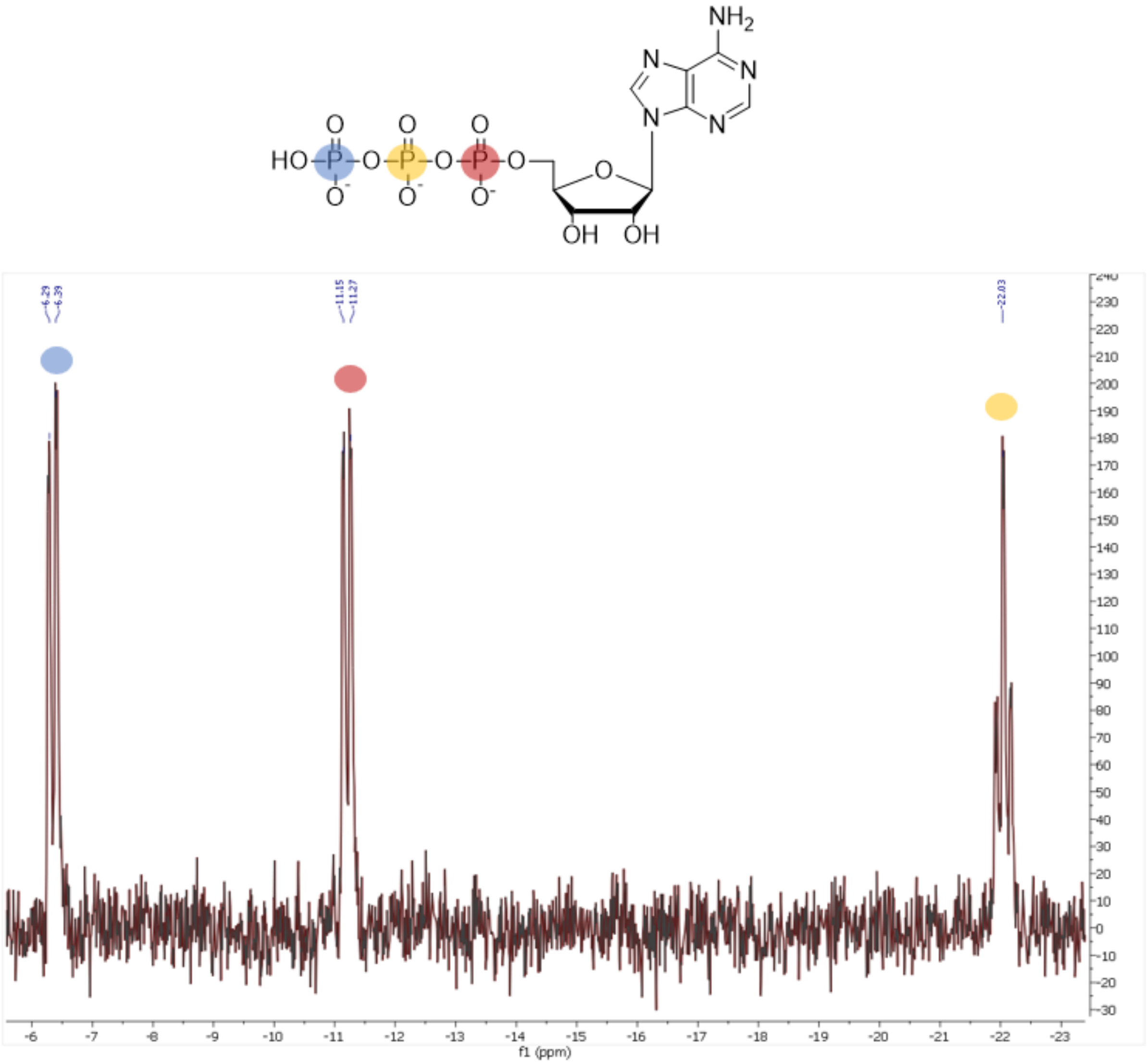


**Figure S24: Overlay of $^{31}P$ NMR spectra of ATP and ATP co-incubated with OD polymer.**

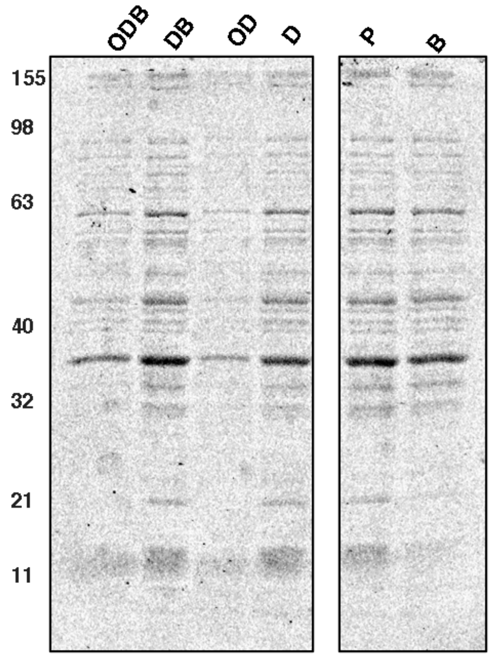


**Figure S25: Coomassie stain of gel in Figure 5A**. MW ladder in kDa.

**Table S4: Summary of assay results for each permeabilization reagent.** Gel intensity values calculated from fluorescence intensity of **B-TMR-ATPγS** labeling total protein stain intensity of HK853. Uptake calculations for PI, NPN and Resazurin assay are located in the methods section.

*E. Coli* Summary Table

| Sample | Gel Intensities | PI Uptake | NPN Uptake | Resazurin Uptake |
|---|---|---|---|---|
| **ODB** | 100% | 5.60 | 0.06 | 1.89 |
| **DB** | 100% | 4.36 | 0.57 | 3.10 |
| **OD** | 94% | 10.80 | 2.88 | 2.52 |
| **D** | 72% | 14.12 | 2.31 | 3.11 |
| **PMBN** | 22% | 10.86 | 1.30 | 1.44 |
| **SNTT** | 37% | 3.00 | 5.48 | 0.72 |

*B. Subtilis* Summary Table

| Sample | PI Uptake | Resazurin Uptake |
|---|---|---|
| **ODB** | 0.54 | 0.64 |
| **DB** | 0.64 | 0.51 |
| **OD** | 1.04 | 0.85 |
| **D** | 1.15 | 0.83 |

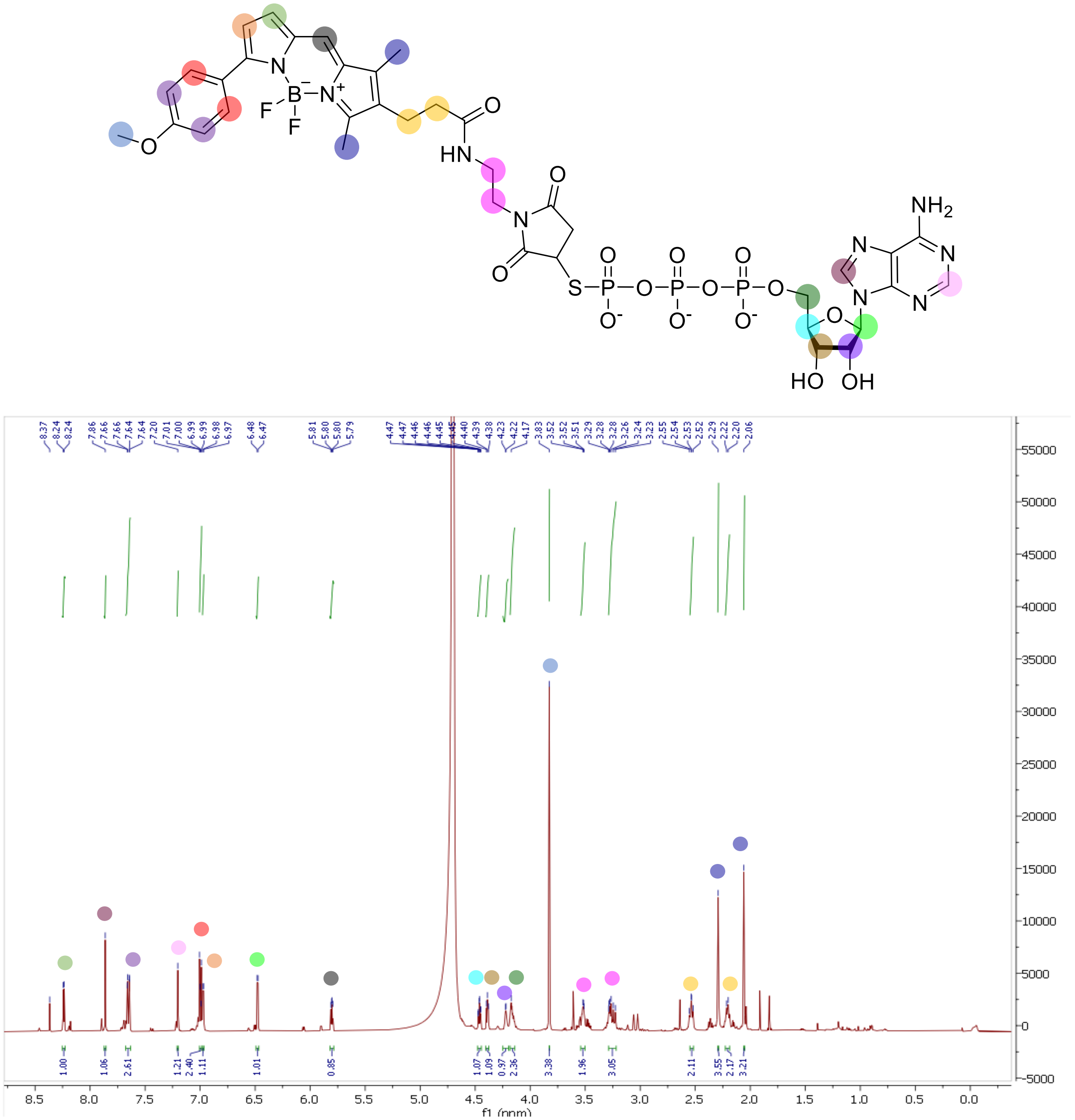

**Figure S26:** $^{1}$H NMR of **B-TMR-ATPγS**.

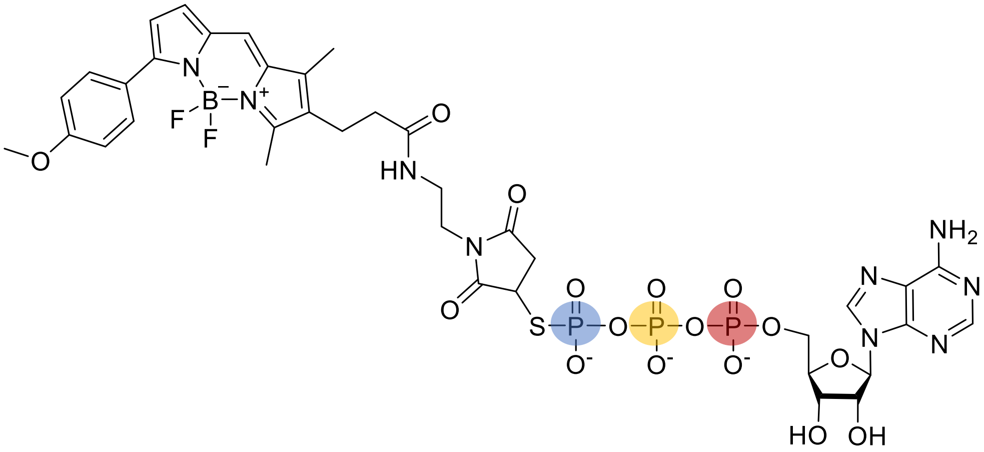


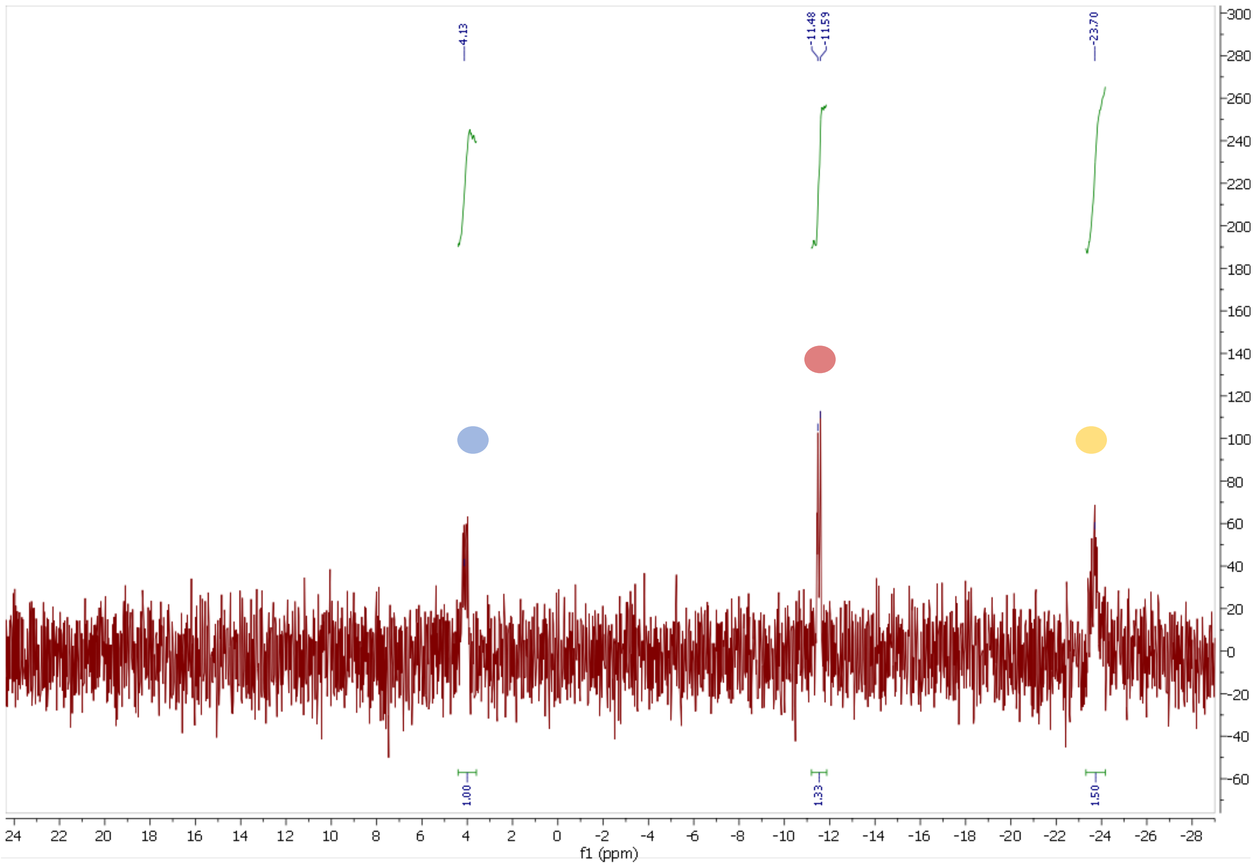


**Figure S27:** $^{31}$P NMR of **B-TMR-ATPγS**.
